# Multi-omic profiling reveals impaired luteal resolution and CHK2-regulated inflammation in ovarian aging

**DOI:** 10.64898/2026.09.30.755521

**Authors:** Liubin Yang, Zhongwei Sun, Xinlong Luo, Carolina Leynes, Isabelle Lorenzo, Ashley Pawelka, Meaghan Donahue, Mancy Tong, Hugh S. Taylor, Yong-hui Jiang, Brendan Lee, Tao P. Wu, Jason D. Heaney

## Abstract

The ovary ages earlier than most other organs, but the molecular mechanisms that drive this decline remain unclear. To gain more insight into this process, we profiled ovaries from young and reproductively aged mice after synchronized ovulation and found that aging was marked by the accumulation of luteal cells with increased steroidogenic, antigen presentation and inflammatory gene expression. DNA methylation was globally preserved in aged ovaries but showed focal age-associated changes at gene regulatory regions. Protein profiling identified an increase in checkpoint kinase 2 (CHK2), a DNA damage response protein encoded by the *CHEK2* gene associated with the timing of natural menopause in humans. CHK2 and DNA damage markers were increased in aged mice and human ovarian somatic cells. Transient CHK2 inhibition in aged mice increased the proportion of mature oocytes and blastocyst formation and reversed inflammatory signatures in granulosa cells. These findings identify defective luteal resolution and inflammation as features of ovarian aging and suggest that CHK2 induces a reversible age-associated inflammatory signature. Our results show that reproductive decline may remain pharmacologically modifiable during ovarian aging via regulation of the inflammatory response.

## Introduction

Ovarian aging shows accelerated decline, often two decades before most other organs^1–3^. This decline in turn leads to infertility, irregular menstrual cycles, and menopause, with significant effects on women’s health. Studying the mechanisms that drive ovarian aging could identify strategies to improve fertility and reproductive longevity.

Mice recapitulate key features of human ovarian aging, including the decline of the ovarian reserve, irregular reproductive cycles, and eventual cessation of ovulation and hormone production^4–6^. Comparative studies indicate that ovaries from 9-12-month-old mice model ovarian aging in humans in their late 30s to mid-40s, and share 20-31% age-related gene-expression changes in oocytes^7^. This age range is particularly relevant to assisted reproductive technology (ART) in which treatment success sharply declines with advancing maternal age from 52% live-birth per intended egg retrieval for patients < 35 years to 3-18% for >38 years^8^.

Growing evidence suggests that the ovarian microenvironment contributes to reproductive aging^3^. Aging ovaries develop increased fibrosis and tissue stiffness^9–11^, and single-cell studies have identified inflammation, oxidative stress, altered immune cells as hallmarks of aging^2,12–15^. Genome-wide association studies of age at natural menopause pinpoint DNA damage response and hematopoietic stem cell pathways^16–18^, but how these pathways relate to age-associated inflammation and tissue dysfunction remains unclear.

Although age-related decline in oocyte number and quality are well recognized features of ovarian aging, the cellular and epigenetic changes that accompany ovarian aging remain poorly understood. Previous ovarian-aging atlases have typically focused on single- or dual-omics approaches in natural cycling or incompletely staged ovaries, potentially making it difficult to distinguish age-dependent changes from cycle-dependent tissue remodeling^19,20^. Our study provides an analysis of single-nucleus transcription, whole-genome DNA methylation, and protein signaling in young and reproductively aged mouse ovaries collected at a matched, gonadotropin-synchronized ovulatory state, with human cell validation and functional studies in mice. We define molecular changes in ovarian aging and identify a potential link between DNA damage response and inflammation.

We show that reproductive aging is associated with the persistence and expansion of inflammatory luteal tissue and increase in checkpoint kinase 2 (CHK2). Single-nucleus analysis showed an expansion of luteal cells characterized by steroidogenesis and JAK–STAT-associated gene signatures and altered ligand–receptor interactions with endothelial cells. Whole-ovary DNA methylation showed focal remodeling at regulatory regions associated with immune related genes and progesterone-mediated oocyte maturation, whereas de novo hypermethylated regions included Msh2, a mediator of meiotic recombination and DNA repair. Proteomic analysis similarly confirmed increased inflammatory signaling, steroid-hormone receptor abundance and an increased DNA-damage sensor CHK2 protein. In addition, pharmacological inhibition of CHK2 during superovulation improved oocyte maturation and blastocyst formation in aged mice and reduced immune-associated gene expression in human granulosa cells.

These findings link defective luteal resolution, inflammation and DNA-damage sensor CHK2 in ovarian aging and provide mechanistic support for human genetic studies that identified loss-of-function variants in *CHEK2* with later age at natural menopause^16,17^, thereby identifying CHK2 as a potentially modifiable regulator of reproductive aging. More broadly, our results suggest that reproductive aging is not driven solely by depletion of the oocyte reserve, but also by potentially modifiable dysfunction within the ovarian somatic environment, providing a framework for developing interventions to preserve fertility and ovarian endocrine function.

## Results

### Aged ovaries show impaired ovulatory response and persistent luteal tissue

Ovarian structure and gene expression vary across estrous cycles and stages of folliculogenesis^19–21^. To reduce potential variation due to differences in cycle stage, we synchronized at least 80% of estrus cycles^22^ by superovulating young (4-5 weeks) and reproductively aged (10-12 months) C57BL6/NJ female mice. After oocyte harvest, we collected ovaries for downstream multi-omic analyses (Fig. 1a). We observed that with age, ovulatory responses declined from 76% (SD 9.8) to 42% (SD 21) (Fig. 1b), and that the proportion of mature oocytes decreased from 63% (SD 19) to 34% (SD 21) (Fig. 1c). On histology, young ovaries exhibited abundant developing and ovulated follicles, whereas aged ovaries had few developing follicles and were largely filled with eosinophilic luteal tissue (Fig. 1d). Periodic Acid-Schiff staining, which highlights carbohydrate-rich zona pellucida of oocytes, further revealed carbohydrate-rich structures within the ovarian medulla and weak staining within the corpora lutea of aged ovaries (Fig. 1e). Thus, synchronized aging ovaries show a reduced ovulatory response, lower proportion of mature oocytes, and extensive residual luteal tissue, consistent with previous studies using synchronized ovulation^23^.

**Fig. 1:**
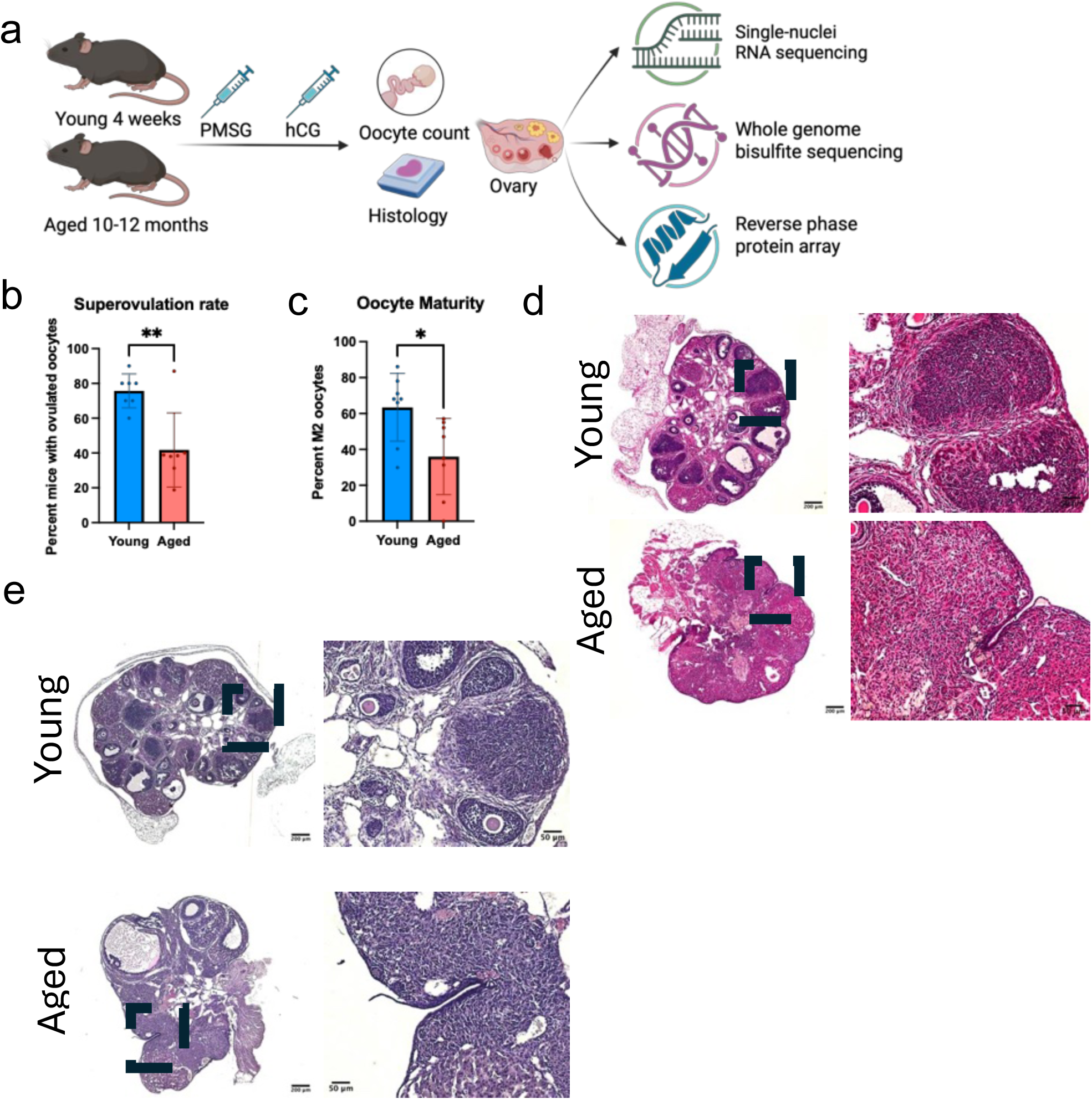
Synchronized superovulation shows declines in ovulatory response and oocyte maturity with abnormal ovarian histology. **a**, Experimental design showing young (4 weeks old) and aged (10 to 12-month-old) mice undergoing superovulation with pregnant mare serum gonadotropin (PMSG) and human chorionic gonadotropin (hCG), oocyte retrieval, and collection of ovaries for histology and multi-omic analyses. **b**, Bar graph of the percentage of mice with oocytes retrieved in each cohort. **c**, Bar graph of the percentage of mature metaphase II (M2) oocytes among all retrieved oocytes. In b and c, each dot represents an independent experimental cohort of young (n=5) and aged (n=5-10) mice. **d**, Hematoxylin and eosin staining of young and aged whole ovaries. **e**, Periodic acid-Schiff staining of young and aged whole ovaries.

### Single nucleus RNA sequencing identifies predominant luteal cells

To understand the changes in cell populations and their gene expression, we performed single-nucleus RNA sequencing of whole ovaries from young and aged mice (4-5 pooled mouse ovaries from 3 independent experiments). We annotated unsupervised clusters using enriched genes and pathways and compared our findings with published ovarian datasets^2,19,20,24–27^ (Extended Data Fig. 1a and Extended Data Tables 1 and 2). We identified fourteen transcriptional clusters, including five corpus luteum (CL) states, four granulosa states, endothelial, theca, stromal, epithelial, and erythrocyte populations (Extended Data Fig. 1b, c, Fig. 2a). Because these clusters were defined transcriptionally, CL and granulosa clusters may represent different cell states rather than distinct cell types. The most significant age-related difference was the expansion of the CL progressing cluster in the aged ovaries (Fig. 2b,c), consistent with our histologic observation of increased luteal tissue.

**Fig. 2:**
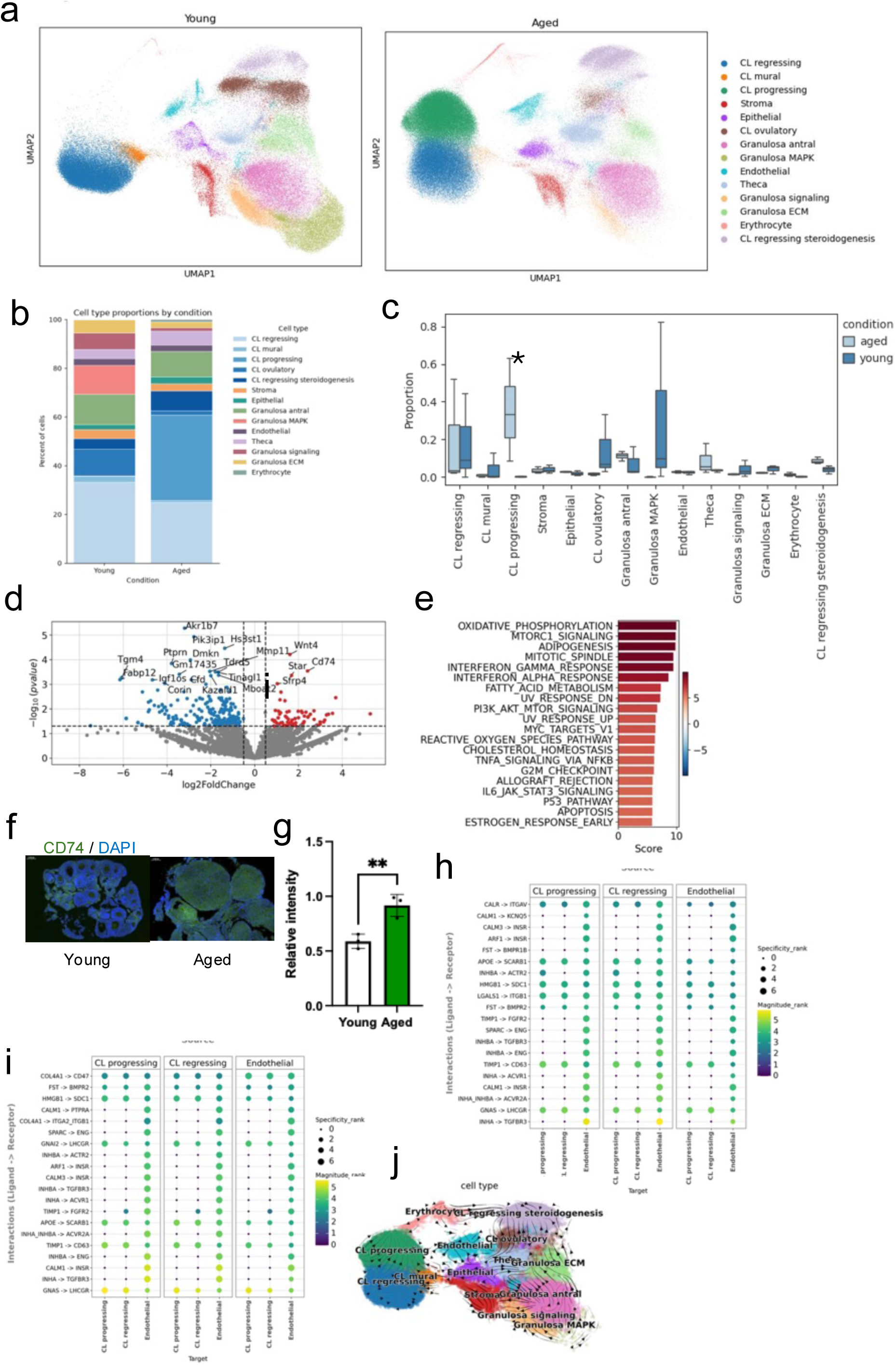
Single-nucleus sequencing reveals an enlarged population of residual corpus luteal cells with upregulated immune-related genes. **a**, Uniform manifold approximation and projection (UMAP) of integrated single-nucleus RNA-sequencing data, colored by annotated clusters in young (left panel) and aged (right panel) groups. **b**, Proportions of annotated cell clusters in young and aged ovaries. **c**, Boxplots of cell-type proportions analyzed using scCODA. The asterisk denotes a credible compositional change at a target false discovery rate of 0.05. **d**, Volcano plot of differentially expressed genes between young and aged mice in the CL progressing cluster, assessed using Wald tests of the pyDESeq2 framework in decoupler. **e**, MSigDB Hallmark gene set activity inferred using a univariate linear model and ranked by t-score. **f**, Immunofluorescent image of CD74 stained young and aged ovaries and **g**, quantification of their relative fluorescent intensity (right panel) using Mann Whitney test over n=2 mice. **h**, Predicted ligand-receptor interactions among CL progressing, CL regressing, and endothelial cells in young ovaries. **i**, Predicted ligand-receptor interactions among CL progressing, CL regressing, and endothelial cells in aged ovaries. Only the top 20 interactions with p-value <0.05 are shown. Dot size represents the specificity rank across all cell type pairs and color represents the consensus interaction magnitude rank. **j,** RNA velocity vectors and the inferred velocity field projected onto a UMAP plot of annotated ovarian cell clusters. Arrows represent RNA velocity vectors.

### Aged luteal cells with steroidogenic, metabolic, and antigen presentation expression

To determine the transcriptomic changes of luteal cells with age, we next studied age-related gene expression in the CL progressing cluster using pseudobulk analysis with biological samples as independent observations. We found that there were 1,996 downregulated and 69 upregulated genes (p<0.05, |log2FC| ≥0.5) (Extended Data Table 3, Fig. 2d). PROGENy^28^ analysis predicted increased JAK-STAT and VEGF pathway activity in aged luteal cells (Extended Data Fig. 1d). Genes contributing to the JAK-STAT pathway included antigen presentation and complement genes *Cd74, C1qb, C1qa, H2-Aa, H2-Ab1* (Extended Data Fig. 1e). There was also an enrichment of oxidative phosphorylation (*Dlat, Prdx3, Suclg1, Acaa2, Bckdha*), and mTORC1 signaling (*Sc5d, Hmgcs1, Idi1, Cyp51, Nampt*), adipogenesis, mitotic spindle, and interferon gamma and alpha pathways (Fig. 2e). Independent batch correction using Harmony likewise identified expanded luteal populations with increased antigen presentation genes (Extended Data Fig. 1f-i), supporting the robustness of these findings. These findings indicate that aged luteal cells acquire increased steroidogenic and metabolic gene expression alongside antigen-presentation and inflammatory signatures, consistent with observations of increased inflammation in aging ovaries^12–14,29^.

We also observed that *HFE,* an iron-homeostasis regulator, was nominated as an upstream regulator driving differential expression changes (Extended Data Fig. 2a,b). Iron is required for heme-containing cytochrome P450 steroidogenic enzymes^30^, and ovaries have one of the highest levels of total *HFE* mRNA of all tissue^31^. Abnormal ovarian iron accumulation has been reported with reproductive aging^32^. Thus, these findings imply that aged luteal cells may upregulate steroidogenesis and iron-overload related stress-metabolic programs.

To validate the antigen-presentation phenotype, we examined the CD74 protein in an independent cohort at another research institution. CD74 expression was increased in luteal cells of aged ovaries (Fig. 2f,g). CD74-positive cells were also present in human ovaries (Extended Data Fig. 2c). Aged mouse ovaries exhibited 7-fold more CD3-positive T cells than young mouse ovaries (Extended Data Fig. 2d), consistent with previous reports of increased T-cell infiltration with age^33^. These findings suggest increased antigen-presentation and immune cell presence with ovarian aging.

### Immune-associated gene expression in extracellular RNA

Because extracellular ambient RNA can influence single-cell analyses, we repeated the analysis without ambient RNA removal. We found that the aged ovaries again showed expansion of the CL population (Extended Data Fig. 2e), with increased expression of MHC class II, Wnt, and inflammatory genes (Extended Data Fig. 2f). Transcription-factor analysis predicted increased activity of immune regulators *Spi1, Sp1,* and sterol regulators *Srebf2* and *Srebf1* (Extended Data Fig. 2g). Gene ontology analysis additionally identified immune function pathways that were absent in our prior analysis with ambient RNA removal: neutrophil degranulation, leukocyte activation, and innate immune response (Extended Data Fig. 2h). IPA analysis predicted an IL-1β-driven inflammatory response (Extended Data Fig. 2i). These data indicate that the aged cohort contained more immune related genes in the extracellular RNA than the young cohort, potentially indicative of increased cellular fragility or inflammation-related damage.

### Aging alters predicted luteal-endothelial communication

Because corpus luteum function depends on vascular remodeling, we examined predicted ligand-receptor interactions between luteal and endothelial cells. Using LIANA+, we ranked predicted ligand-receptor interactions using consensus magnitude rank, which integrates multiple interaction-strength metrics across cell-cell communication methods. We identified several predicted ligand-receptor interactions shared between young and aged ovaries, including those involved in cholesterol handling and extracellular matrix signaling, such as APOE-SCARB1 and HMGB1-SDC1. Young ovaries showed predicted signaling involving activin/inhibin TGF-β family signaling (INHA-TGFBR3, INHBA-ENG, INHA-ACVR2A, FST-BMPR1B), consistent with luteal paracrine signaling; and endothelial signaling (SPARC-ENG, TIMP1-CD63/FGFR2, LGALS1-ITGB1, CALR-ITGAV) (Fig. 2h). In aged ovaries, highly ranked interactions instead shifted toward extracellular matrix and cell-adhesion signaling, particularly collagen IV interactions with integrin receptors, including COL4A1-ITGA2, COL4A1-ITGB1, as well as collagen IV with anti-phagocytosis COL4A1-CD47 (Fig. 2i). These predictions suggest that aging shifts luteal-endothelial communication towards collagen- and extracellular matrix programs.

### Regressing corpus luteum trajectory

To infer relationships among ovarian cell states, we performed RNA velocity analysis. The inferred velocity field connected corpus luteal (CL) progressing cluster with the CL regressing population, with directional flow toward the regressing state, likely consistent with progression from an active luteal state towards luteal regression (Fig. 2j). Granulosa cell clusters followed a separate trajectory from antral to luteinizing and ovulatory states (Extended Data Fig. 2j-l). PHATE analysis^34^ also showed that luteal populations occupied a connected transcriptional manifold, consistent with states spanning luteinization, remodeling, and regression (Extended Data Fig. 2m). Granulosa populations similarly connected toward luteal states, whereas stromal, theca, endothelial, and epithelial populations remained more distinct, reflective of divergent lineages. Together with the accumulation of CL progressing cells in aged ovaries, these analyses are consistent with altered or incomplete resolution of the corpus luteum during reproductive aging.

### Preserved global DNA methylation

DNA methylation is a major hallmark of aging with tissue-specific aging signatures^35^, but genome-wide studies of ovarian aging remain limited^36–38^. We therefore performed whole genome bisulfite sequencing on ovaries collected after synchronized ovulation, yielding approximately 252 million reads per sample, with 81-83% mapping rates and 15-27x mean genomic coverage (Extended Data Table 4). Our findings showed that global methylation distributions were broadly similar between young and aged ovaries across CG, CHG, and CHH methylation (Fig. 3a) and across major genomic annotations, including CpG islands and shores, promoters, 5’ UTRs, exons, introns, 3’ UTRs, and repetitive elements (Extended Data Fig. 3a, b). Both groups also showed the expected depletion of methylation around transcription start sites and higher methylation across gene bodies (Extended Data Fig. 3c). Age-related gains and losses of methylation were distributed across chromosomes (Extended Data Fig. 3d). Thus, global DNA methylation architecture remains largely preserved with ovarian aging, consistent with that observed in porcine and mouse ovarian studies^36,37^.

**Fig. 3:**
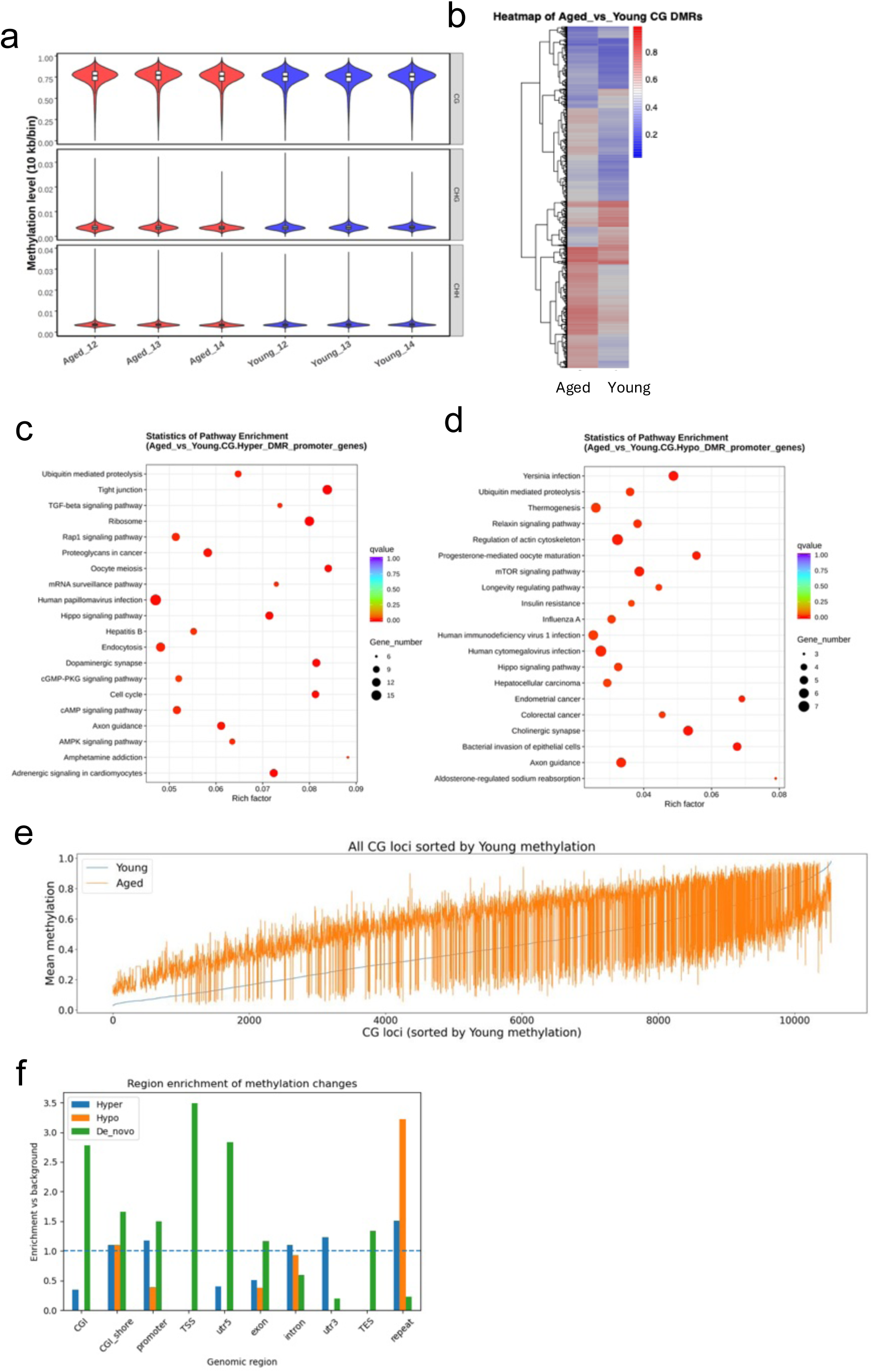
Whole genome bisulfite sequencing shows site-specific hypo- and hypermethylation and de novo methylation at regulatory sites. **a**, Violin plot of methylation level over 10kb bins for CG, CHG, and CHH contexts in individual young and aged samples. **b**, Heatmap of mean methylation levels in differentially methylated CG regions in aged and young ovaries. **c**, KEGG pathway enrichment for genes associated with hypermethylated promoter regions. **d**, KEGG pathway enrichment for genes associated with hypomethylated promoter regions. In c and d, the horizontal axis shows the rich factor, color denotes the q value, and dot size denotes the number of genes. **e**, Mean methylation levels at CG loci in young (blue) and aged (orange) ovaries, sorted by methylation level in the young group. **f**, Fold enrichment of hypermethylated, hypomethylated, and age-acquired methylated regions across genomic annotations relative to genomic background. Hyper and Hypo denote methylation increases and decreases greater than 0.25, respectively. De novo denotes regions with 0–5% methylation in young ovaries and >5% methylation in aged ovaries. The dashed line indicates an enrichment ratio of 1; values >1 indicate overrepresentation and values <1 indicate depletion. CGI, CpG island; TSS, transcription start site; utr5, 5′ untranslated region; utr3, 3′ untranslated region; TES, transcription end site.

### Focal hypermethylation at regulatory regions

To identify locus-specific changes in methylation, we performed differential methylation analysis and found 6,514 CG-context differentially methylated regions (DMRs), including 4,959 hypermethylated and 1,555 hypomethylated regions, of which 3,795 were associated with annotated genes (Extended Data Table 5, Extended Data Fig. 3e). Thus, age-associated methylation change showed more hypermethylated sites, similar to observations in postmenopausal human ovaries^38^. DMRs included both gains and losses of methylation (Extended Data Fig. 3f, Fig. 3b). Many occurred in regions with low baseline methylation and regions with intermediate-to-high methylation levels of 0.6-0.85 (Extended Data Fig. 3g). Hypermethylated CG DMRs were enriched at CpG islands, CpG island shores, promoter, and 5’ untranslated regions (Extended Data Fig. 3h) compared to CHG and CHH sites (Extended Data Fig. 3i and j), indicating age-associated methylation changes at focal gene-regulatory regions. Similar to the results seen in our young mice, age-related methylation gains at previously unmethylated (<5%) sites are also seen in young humans, mice, and monkeys^39^.

### Focal differential methylation

Within the DMRs, we next examined the distribution of age-associated methylation changes across gene bodies, promoters, and regions. We found that hypermethylated gene bodies were enriched for intracellular signal transduction (*Rcan1, Arhgef3, Rapgef2*) and small GTPase signaling (*Sipa1l2, Sh2d3c, Arhgef19*) (Extended Data Fig. 4a). KEGG analysis identified pathways in metabolism (*Haao, Gm13443, Synj2*), axon guidance (*Ntn1, Sema4a, Camk2d*), Rap1 (*Vav3, Src, Fgf10)*, and MAPK signaling (Extended Data Fig. 4b). Hypomethylated gene bodies were also enriched for MAPK signaling (*Rras2, Nf1, Fgf10*), together with proteoglycans (*Frs2, Elk1, Drosha*) and Wnt signaling pathways (*Nfatc2, Wnt6, Nfatc1*) (Extended Data Fig. 4c). Thus, although individual loci gained or lost methylation with age, both sets of loci share signaling and tissue-remodeling pathways. Several of these pathway categories, including axon guidance, proteoglycans in cancer have also been reported in methylation studies of multiple human tissue types in aging^38^. Additionally, Rap1, axon guidance, focal adhesion, and MAPK signaling pathways have been reported in methylation studies of postmenopausal ovaries^38^. Overall, these results indicate that some methylation changes are shared across other human tissues while others may be ovary specific such as MAPK signaling.

We next asked whether age-associated methylation changes also affected promoter regions, which have a direct role in transcriptional regulation. Promoter hypermethylation was enriched in regions associated with phosphatase regulator activity (*Ppp2r5d, Ppp1r2-ps4, Gm13169*) and DNA binding (*Kdm2b, Runx2, Irx2*) (Extended Data Fig. 4d) and in pathways related to tight junctions (*Prkag1, Synpo, AC161165.4*), ribosome (*Gm45497, Rps4l, Gm15846, Rpl36a-ps3*), dopaminergic synapse signaling (*Gm13244, Creb3l4, Gm26794*), oocyte meiosis (*Ywhab, Gm13244, Gm15045*), and cell cycle (*Ywhab, Gm15045, Cdkn1c, Anapc7*) (Fig. 3c). In contrast, hypomethylated promoter-associated DMRs were enriched for progesterone-mediated oocyte maturation (*Rps6ka3, Adcy6, Gm12992, Pik3cd, Rps6ka6*) and mTOR signaling (*Sgk1, Eif4ebp1, Rps6ka6*) (Fig. 3d). Together, these findings indicate that age-associated promoter methylation changes mapped to pathways involved in cellular signaling, cell-cycle regulation, and ovarian function.

We further identified 232 age-acquired methylated regions, defined as loci with <5% methylation in young ovaries that have >5% methylation in aged ovaries (Fig. 3e). Relative to genomic background, these regions were preferentially localized to transcription start sites, 5’ UTRs, and CpG islands (Fig. 3f). Associated genes included the developmental regulators *Foxf1, Foxp4*, *Zic1,* and *Zic4* and the DNA mismatch repair gene *Msh2* (Extended Data Fig. 4e). Together, these findings indicate that ovarian aging is associated with focal methylation gains at gene regulatory regions, including loci involved in development and DNA repair.

### Proteomic profiling

Because protein abundance in the ovary cannot always be inferred from RNA expression^40^, we performed reverse phase protein array profiling^41^ of synchronized post-ovulatory ovaries of young and aged mice (Fig. 4a). Unsupervised clustering identified age-associated differences in protein abundance (Fig. 4b, Extended Data Table 6). Aged ovaries showed an increase in hormone response (PGR, AR), cell signaling (STAT4, IGF1R, MAPK1), and checkpoint kinase 2 (CHK2) proteins. They also displayed decreased abundance of proteins involved in epigenetic regulation (DNMT1, EZH2), DNA synthesis and repair (RRM2), cyclooxygenase (PTGS2) activity, and ERBB2 signaling, as well as lower levels of the mTOR signaling protein RPS6KB1 than young ovaries (Fig. 4c). Pathway analysis predicted activation in inflammatory IL-12, p53, and aryl hydrocarbon receptor signaling and inhibition of ovarian folliculogenesis and pulmonary fibrosis genes (Fig. 4d). Gene set enrichment confirmed that proteins were enriched for ovary bulk tissue (Extended Data Fig. 5a). We also observed predicted activation of apoptosis as well as inhibition of cell growth and viability in aged ovaries compared to young ovaries (Extended Data Fig. 5b).

**Fig. 4:**
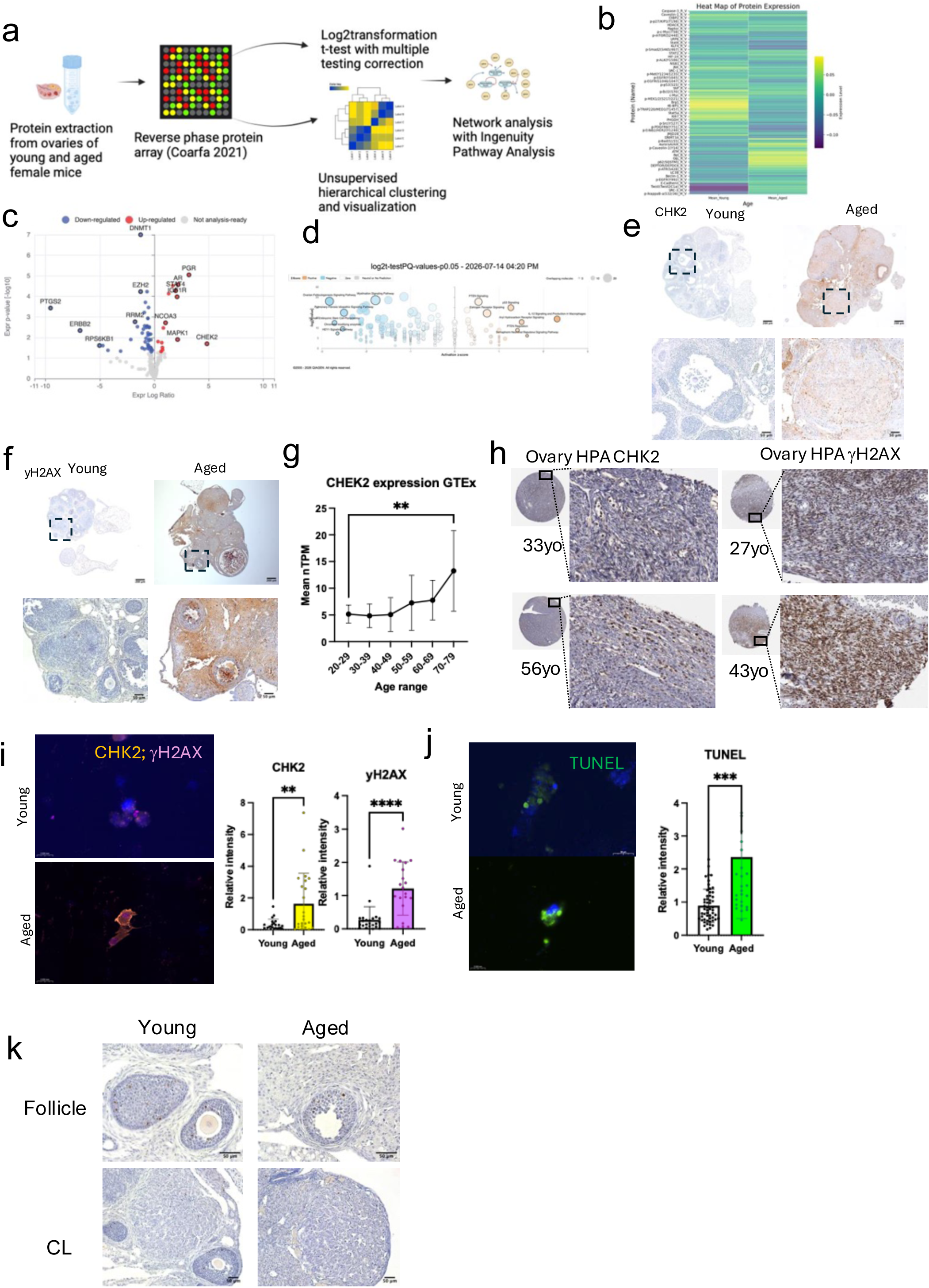
Proteomics show high levels of inflammation and DNA damage in somatic cells. **a**, Illustration of the collection of pooled ovaries from young (4 weeks old) and aged (10–12-month-old) mice (n=5 for each group) antibody based reverse phase protein array of 261 validated antibodies, statistical analysis with normalization, unsupervised hierarchical clustering, and network analysis. **b**, Heatmap of mean normalized protein expression in young and aged groups. **c**, Volcano plot of differentially expressed proteins between young and aged groups. Color indicates expression change. Ingenuity Pathway Analysis software mapped mouse data to human genes. **d**, Bubble plot of predicted activated and inhibited canonical pathways. Colors indicate Z-score. Annotated pathways stated on plot. **e**, Immunohistochemistry of representative whole ovary section of young and aged samples stained with anti-CHK2 antibody. **f**, Immunohistochemistry of representative whole ovary section of young and aged samples stained with anti-gH2AX antibody. **g**, Line plot of mean normalized transcripts per million (TPM) of *CHEK2* in human ovaries, derived from a gene model with isoforms collapsed to a single gene by GTEx database, across donor age groups. Error bars indicate standard deviation. **h**, Histological images of human ovaries stained with anti-CHK2 antibody (left panels) and anti-gH2AX antibody (right panels) obtained from Human Protein Atlas (HPA). yo is years old. **i**, Immunofluorescent co-staining of primary human lutein granulosa cells with anti-CHK2 and anti-gH2AX antibodies derived from representative donor per group. Experiment independently repeated 3 times. Right graph is quantification of fluorescent intensity normalized to DAPI. **p<0.01, ****p<0.0001. **j**, Immunofluorescent TUNEL staining of primary human lutein granulosa cells. Right graph is quantification of fluorescent intensity. ***p<0.001. **k**, Histological images of representative aged and young ovaries stained with anti-phospho-CHK2 Thr68 antibody for follicle (top panels) and corpus lutea (lower panels).

Among the upregulated proteins in aged ovaries was CHK2, a DNA damage response kinase encoded by the *CHEK2* gene, whose loss-of-function variants are associated with age at natural menopause and ovarian reserve^16,42,43^. *CHEK2* is known to regulate oocyte integrity^44,45^. In our study, immunostaining revealed increased CHK2 in somatic cells of aged mouse ovaries (Fig. 4e), suggesting that CHK2 may play a role in somatic DNA damage response in addition to its established role in oocyte quality control.

### CHK2 and DNA damage markers

We next examined markers of DNA damage and repair using immunohistochemistry. We determined that aged mouse ovaries showed increased γH2AX, a marker of double stranded DNA breaks in stromal and cumulus cells (Fig. 4f). We also found that TUNEL staining was weak in luteal and stromal cells, indicating potential nonspecific staining (Extended Data Fig. 5c). Markers of homologous recombination and non-homologous end joining showed no clear age-associated differences (Extended Data Fig. 5c). Thus, aged ovarian somatic cells showed increased DNA-damage signaling without a corresponding increase in the repair markers examined.

In human ovary samples, we found that *CHEK2* is highly expressed in the Genotype-Tissue Expression (GTEx) dataset (Extended Data Fig. 5d) and that *CHEK2* expression increased with age, with higher expression in donors aged 70-79 years than in those aged 20-29 years (Fig. 4g). Likewise, CHK2 and γH2AX were also present in human ovaries in the Human Protein Atlas (Fig. 4h). Granulosa cells from women aged ≥40 years undergoing IVF showed higher CHK2 and γH2AX levels than women aged <35 years (Fig. 4i), as well as increased TUNEL staining (Fig. 4j). These findings support increased CHK2 expression and DNA-damage in aged human ovarian somatic cells, consistent with other studies^46^.

To examine downstream DNA-damage pathways, we surveyed expression changes of downstream members by immunohistochemistry and the above single-nuclei RNA sequencing data. Embryos in the blastocyst stage derived from aged mouse oocytes showed increased p53 expression (Extended Data Fig. 5e), whereas transcriptomic analyses of CL progressing cluster did not identify increased DNA damage sensing pathways such as cGAS-STING, ATM, or NF-kB signaling pathways (Extended Data Fig. 5f). Despite increased total CHK2, canonical phosphorylation of CHK2 at the Thr68 site by ATM kinase showed little age-related difference in mouse ovaries compared to young (Fig. 4k). These findings suggest that increased CHK2 in aged mouse ovaries is not accompanied by changes in canonical ATM phosphorylation site. The findings are consistent with, but do not yet prove, a role for elevated CHK2 outside of the canonical DNA damage response in aging.

### Transient CHK2 inhibition improved oocyte maturation

Human genetic studies implicate *CHEK2* in influencing reproductive lifespan. Specifically, germline *Chek2* deletion delayed ovarian aging in mice^16^. We therefore tested whether inhibition of CHK2 during reproductive aging could improve ovarian function. For this purpose, we treated reproductively aged mice with a potent CHK2 inhibitor CCT241533 during ovarian stimulation (Fig. 5a). We found an increase in the proportion of mature oocytes (Fig. 5b). An independent cohort of aged mice from the National Institute on Aging colony also showed an increase that was not statistically significant (Extended Data Fig. 6a). To determine whether the response was conserved across genetic backgrounds, we additionally tested aged outbred CD-1 mice and again observed an increase in the proportion of mature oocytes (Fig. 5c). In the NIA C57BL/6 cohort, CHK2 inhibition also increased the number of blastocysts approximately twofold (Fig. 5d). Thus, transient CHK2 inhibition improved selected IVF outcomes in multiple cohorts and mouse strains.

**Fig. 5:**
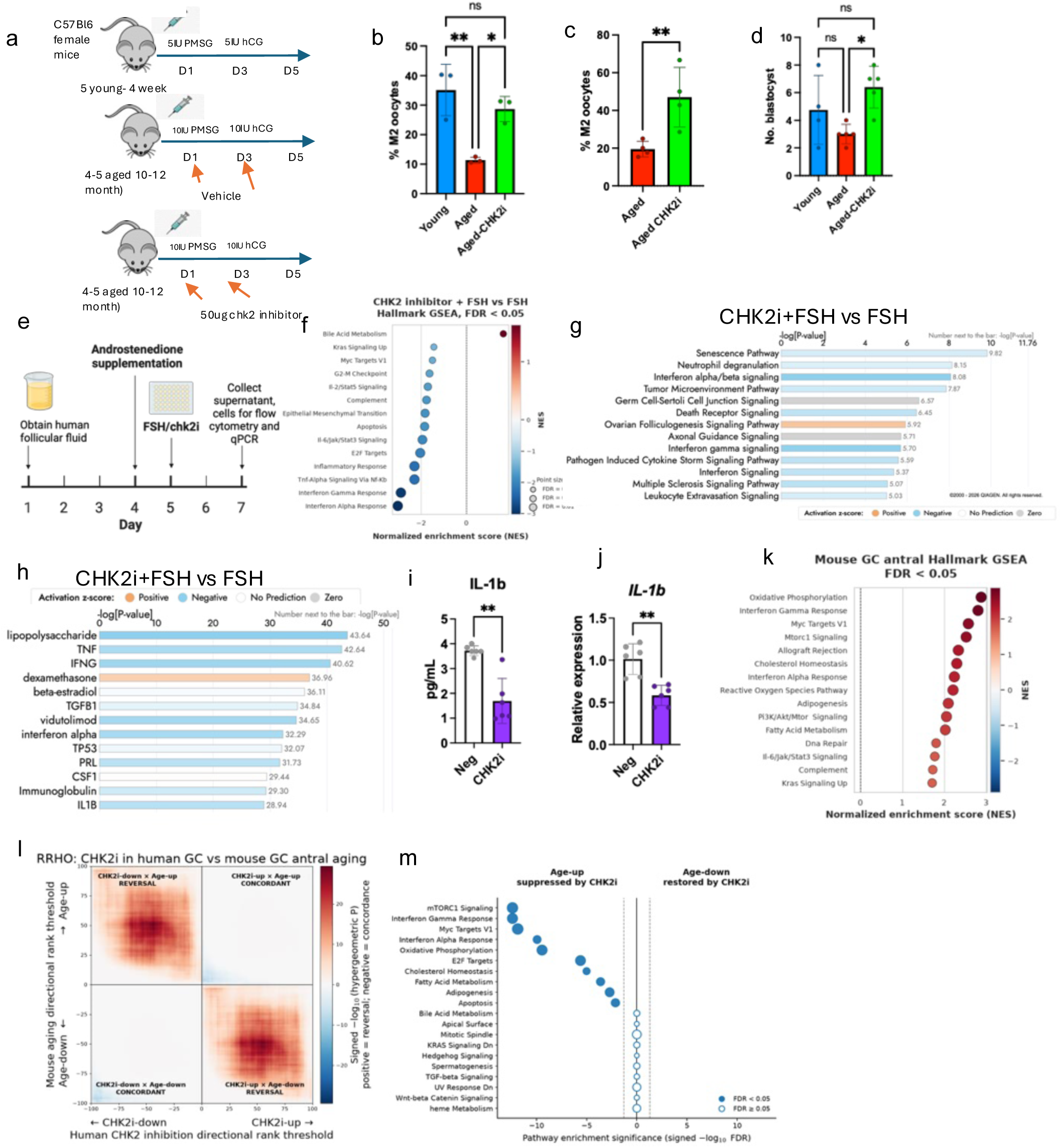
Short term CHK2 inhibition improves oocyte quality and reverses age-related inflammatory genes. **a**, Diagram of superovulation treatment schedule for young (4 weeks old) and aged (10-12 month old) mice (n=5 for each age group). Arrows indicate intraperitoneal injection schedule of pregnant mare serum gonadotropin (PMSG), human chorionic gonadotropin (hCG), and CHK2 inhibitor CCT241533 (CHK2i). **b**, Bar graph of percentage of mature metaphase II (M2) oocytes after superovulation in C57Bl6/NJ strain from Baylor Knockout Mouse Phenotype core. Aged-CHK2i denotes aged mice treated with CCT241533. Bars indicate means with standard deviation. Dots represent experimental cohorts. n=4-6 animals per cohort over two independent experiments. **c**, Bar graph of percentage of M2 oocytes after superovulation in 11-month-old CD-1 mice. Bars show means with standard deviation. Aged-chk2i are aged mice treated with CHK2 inhibitor. Dots represent each mouse. n=4-6. **d**, Bar chart of number of embryos reaching the blastocyst stage in the National Institute of Aging Rodent Colony C57BL/6 cohort. Bars indicate standard deviation. Dots represent individual mice. **e**, Experimental timeline of primary human cell culture, media supplementation, and follicle stimulation hormone (FSH) and CCT241533 (CHK2i) treatment, and sample collection. **f**, Dot plot of MSigDB Hallmark gene sets enrichment comparing CHK2i+FSH versus FSH treated primary human granulosa cells. Pathways with FDR <0.05 are shown. Position and color indicate normalized enrichment score (NES), and dot size represents false discovery rate (FDR). **g**, Bar graph of canonical pathways associated with differentially expressed genes between CHK2i+FSH versus FSH treated primary human granulosa cells. Bar length indicates −log10(P), color denotes predicted pathway activation (orange), inhibition (blue), or no directional prediction (gray) based on the IPA activation z-score. **h**, Bar graph of predicted upstream regulators based on differentially expressed genes between CHK2i+FSH vs FSH treated primary human granulosa cells. Bar length indicates −log10(P), color denotes predicted pathway activation (orange), inhibition (blue), or no directional prediction (gray) based on the IPA activation z-score. **i**, Bar graph of secreted IL-1β concentration in primary granulosa cell culture supernatant after CCT241533 (CHK2i) treatment versus the negative-control condition. **j**, Bar graph of relative expression of IL1B for the same two treatment labels. **p<0.01. **k**, Dot plot of significantly enriched MSigDB Hallmark gene sets in aged versus young mouse granulosa antral cluster (FDR <0.05); position and color indicate NES. **l**, Heatmap of rank-rank hypergeometric overlap (RRHO) analysis of ortholog-matched genes from human granulosa cells treated with CHK2 inhibitor plus FSH vs FSH alone and in aged versus young mouse antral granulosa cell cluster. The horizontal axis shows directional rank thresholds for the human CHK2i response, vertical axis shows directional rank thresholds for the mouse aging response. Color shows signed −log10 hypergeometric *P*, with positive values denoting discordance and negative values denoting concordance. **m**, Dot plot of over-representation analysis of MSigDB Hallmark pathways among genes with opposing aging and CHK2i responses, using RRHO ortholog-matched gene universe as background. Left side shows pathways for genes decreased by CHK2 inhibition but increased with aging. Right side shows pathways for genes increased by CHK2 inhibition but decreased with aging. Horizontal axis shows signed −log10(FDR) with negative values assigned to age-up/CHK2inhibition decreased and vice versa for visualization. Dot size represents the number of overlapping genes. Filled circles indicate pathways enriched at FDR <0.05, open circles indicate FDR >=0.05. For all statistical tests, ns is non-significant, *p<0.05, **p<0.01, ***p<0.001, ****p<0.0001.

### CHK2 inhibition dampens inflammation

To investigate how CHK2 inhibition improves ovarian response, we profiled the protein from ovaries of reproductively aged mice treated with CCT241533 and compared our results to those of untreated mice. We found that the treated mouse ovaries exhibited predicted inhibition of ribonucleotide reductase, HER2 signaling, and fibrosis pathways compared to the untreated ones (Extended Data Fig. 6b). These findings suggest that the effects of CHK2 inhibition may extend beyond canonical DNA-damage signaling to pathways involved in growth-factor signaling and fibrosis in the ovary.

Because cumulus and ovulatory granulosa cells express innate immune programs^47,48^, we evaluated whether CHK2 regulates immune programs by treating human granulosa cells with CHK2 inhibitor CCT241533 with and without follicle stimulating hormone (FSH) (Fig. 5e). We first confirmed that the cultured cells retained functional granulosa-cell characteristics by demonstrating FSH-responsive estradiol production (Extended Data Fig. 6c). Then we conducted bulk RNA sequencing analysis, which revealed that CHK2 inhibition reduced interferon alpha, gamma response, Tnf-α and inflammatory response pathways (p<0.05 for Hallmark enrichment, Fig. 5f, Extended Data Fig. 6d, Extended Data Table 7), with predicted inhibition of senescence pathway, neutrophil degranulation, and interferon alpha and beta signaling by IPA (Fig. 5g, the inhibitor-alone comparison is shown separately in Extended data Fig. 6e). We found predicted inhibition of estradiol, lipopolysaccharide, and IL1B signaling in upstream regulator analysis (Fig. 5h), with inhibitor-alone comparison (Extended Data Fig. 6f). Similarly, IL-1β expression and secretion decreased with CHK2 inhibition (Fig. 5i,j). These results indicate that CHK2 contributes to inflammatory signaling in human granulosa cells.

To determine whether inhibition of CHK2 alters gene expression programs associated with ovarian aging, we compared transcriptional changes induced by CHK2 inhibition in human granulosa cells with age-associated changes in antral granulosa cell cluster from single-cell RNA sequencing of young and aged mouse ovaries. We found several pathways altered with aging in the mouse ovary antral granulosa cell cluster, including oxidative phosphorylation, interferon-γ and interferon-α, and Myc target pathways (Fig. 5k). To assess for any relationship across the two transcriptomes, we then compared ranked gene-expression changes in these two datasets using rank-rank hypergeometric overlap analysis^49^. We observed that genes upregulated by CHK2 inhibition preferentially overlapped with genes downregulated with aging (maximum −log10 hypergeometric P = 28.79), while genes downregulated by CHK2 inhibition preferentially overlapped with genes upregulated with aging (maximum −log10 hypergeometric P = 29.38). In contrast, concordant CHK2 inhibition-up/age-up and CHK2 inhibition-down/age-down overlaps were substantially weaker (maximum −log10 hypergeometric P = 9.01 and 8.60, respectively, Fig. 5l). These results indicate that CHK2 inhibition drives transcriptomic changes that oppose some of those of aging.

To identify biological pathways underlying the transcriptome changes, we performed over-representation analysis. Our data show that genes that were upregulated with aging but downregulated following CHK2 inhibition were significantly enriched for inflammatory, stress-response, transcriptional, and metabolic programs: *Mtorc1* signaling, interferon-γ and interferon-α response, *Myc* targets, and oxidative phosphorylation pathways (Fig. 5m). In contrast, genes downregulated with aging that increased after CHK2 inhibition did not show significant enrichment for Hallmark pathways but were enriched for transcription-related biological processes (Extended Data Fig. 6g). Together, these findings indicate that CHK2 inhibition in human granulosa cells induces gene expression changes that broadly oppose age-associated transcriptional changes in mouse granulosa cell clusters, with the strongest reversals in inflammation, stress-response, and metabolic programs.

## Discussion

Our work demonstrates that synchronization of the complex estrous cycle in mice can reveal insights into ovarian aging. Like other mouse studies, our model also demonstrated age-related depletion of follicles and poor oocyte quality and increased residual corpus-luteum-like cells on histology^21,26^. We used synchronized ovulation and transcriptomic, methylome, and proteomic analyses to define molecular features of ovarian aging. Aged ovaries accumulated luteal cells with increased steroidogenic, metabolic mTOR signaling, and antigen presentation programs, and as well as increased CHK2 and markers of DNA damage.

A prominent finding was the accumulation of luteal cells in aged ovaries, resonating with histological observations of luteinized or post-ovulatory steroidogenic stromal cells filling the aged human ovaries^50^. Our single nucleus data provide a molecular correlate of this phenotype. RNA velocity and trajectory analyses placed the expanded aged luteal population along a continuum of luteal remodeling and regression, consistent with impaired resolution of the corpus luteum. These luteal cells showed increased oxidative phosphorylation, steroidogenesis, and antigen presentation programs, including increased expression of *Cd74* and MHC II genes. Similar increases in hormone biosynthesis, inflammation, oxygen-related proteins have also been reported in reproductive aged mouse ovaries^40,51^.

Reproductive aged luteal cells also showed increased antigen presentation and innate immune-related genes. As granulosa and luteinized granulosa cells can intrinsically express immune-related programs^47,48,52^, our data suggest that these signatures may reflect age-related granulosa-intrinsic changes and could be coupled to changes in proportions of immune cells^33^. Increased CD74 expression in aged luteal cells, together with increased T-cell abundance and broader inflammatory signatures, supports coordinated changes both in ovarian somatic cells and their local immune microenvironment. Further studies will be needed to assess the role of the innate immune responses in ovarian aging.

Whole genome bisulfite sequencing revealed that global DNA methylation was largely preserved between young and aged ovaries, with numerous site-specific changes in gene-regulatory regions, predominantly hypermethylation. Compared to young ovaries, aged ovaries also showed focal methylation changes in CG-rich regulatory regions of the genome that correspond to immune-related, cell signaling, mTOR, and DNA repair. Similarly, in other tissues, physiologic aging is associated with differentially methylated sites^38^. Others have also found both hypermethylated and hypomethylated sites in aged human lutein granulosa cells using MethylCap-seq and reduced representation bisulfite sequencing^53^. Additionally, because our analyses were performed on whole ovaries, observed differences between the two may also reflect age-associated shifts in cellular composition, such as increased luteal populations. In future studies, cell-type resolved epigenomic studies will be required to distinguish cell-specific versus cellular compositional changes.

Our protein analyses confirmed an increase in inflammation and identified CHK2 as a potential mediator of age-associated ovarian decline. *CHEK2* variants are associated with later age at natural menopause^16,17^ and CHK2 has an established role in eliminating oocytes with DNA damage^44,45^. Our findings extend this biology to ovarian somatic cells: aged mouse and human ovarian cells showed increased CHK2 with increased γH2AX. However, in aged mouse ovaries, we did not observe a corresponding increase in CHK2 Thr68 phosphorylation or other canonical DNA-repair markers, suggesting that CHK2 increase is not associated with increased activation of the canonical DNA-damage response. Other studies also support our findings that granulosa and stromal cells accumulate DNA double strand breaks with poor DNA repair response^46,54^.

Importantly, transient CHK2 inhibition during ovarian stimulation improved IVF stimulation outcomes, results reproduced at other institutions and for other mouse strains. These experiments suggest that at least some age-associated ovarian dysfunction remains pharmacologically modifiable. However, additional studies are needed to define other key metrics of ovarian aging, pinpoint relevant ovarian cell types, and establish the safety and optimal timing of CHK2 inhibition before considering translational applications.

Emerging studies indicate that outside of its canonical role in DNA damage signaling, CHK2 also regulates inflammatory and immune pathways^55–59^. Consistent with these observations, we found that CHK2 inhibition decreased expression of genes related to fibrosis and inflammation in human granulosa cells. These transcriptional changes opposed some of the age-associated changes in mouse granulosa cell clusters identified by single-nucleus RNA sequencing. These findings in somatic ovarian cells of humans and mice provide a potential mechanistic link between two groups of genes implicated by human studies of age at natural menopause: DNA-damage response and hematopoietic or immune pathways^16,17^. The precise mechanisms linking CHK2 to ovarian immune signaling remain to be unraveled.

Our study has several limitations. Controlled ovarian hyperstimulation is not a natural physiologic state and may amplify age-associated differences^60^. Our analyses also capture a single post-ovulatory time point and therefore may not represent changes across the reproductive cycle. In addition, single-nucleus sequencing limited our ability to detect oocytes and immune cells, likely due to extraction techniques and cell abundances^2^. The human granulosa-cell bulk RNA sequencing experiment also used replicates from a single donor. These are within-donor culture replicates and while it can reduce within-donor variation, the results do not establish reproducibility across donors. Future studies using additional samples, cell-type resolved and longitudinal approaches across synchronized cycle stages will be needed to define how these molecular changes evolve with aging.

Together, our findings identify impaired luteal resolution, increased inflammation, focal epigenetic changes and increased CHK2 signaling as key features of ovarian aging. They further nominate CHK2 as a regulator of ovarian inflammation and declining reproductive function.

## Methods

### Superovulation and collection of mouse oocytes and ovaries

Rodent work was conducted in accordance with the Institutional Animal Care and Use Committee standards and guidelines at Baylor College of Medicine and Yale School of Medicine. At Baylor College of Medicine, C57BL6/NJ mice were housed in a temperature-controlled environment with equal 12 hours of light and dark cycles. Aged mice were retired breeders donated from the Baylor Genetically Engineered Rodent Models Core facility with zero to 3 litters each^61^. At Yale School of Medicine, mice were housed in a pathogen-free facility in a 12-hour light and dark cycle at a temperature of 20-22°C and humidity of 30-70%. Young 4–5-week-old C57BL/6J female mice were obtained from Jackson Laboratory. Aged virginal C57BL/6J female mice were obtained from the Aged Rodent Colony, maintained by Charles River Laboratories from the National Institute on Aging. Female, 11-month-old CD-1 mice were retired breeders donated from the Yale Genome Editing Center.

Young and aged female mice were stimulated with 5IU or 10IU of pregnant mare serum gonadotropins followed by 5IU or 10IU of human chorionic gonadotropin (Sigma CG10-1VL) after 48h by intraperitoneal injection. For CHK2 inhibition treatment, mice were injected with CCT241533 (50 μg per mouse; 1.67mg/kg) or vehicle (5% dimethyl sulfoxide, 95% β-cyclodextrin) (MedChemExpress) via intraperitoneal injection on days 1 and 3 concurrent with PMSG and hCG administration, respectively. Mice were sacrificed and their ovaries and oviducts were harvested for oocyte collection 16-18h later as previously described^61^. The cumulus oocyte complexes were inseminated with fresh sperm from a single 4–5-month-old male C57BL6/NJ donor. Briefly, motile sperm were harvested from cauda epididymis and incubated in methyl-b-cyclodextrin with modified krebs-Ringer bicarb solution (THY) at Baylor or FertiUP (Cosmobio) at Yale. Following conventional insemination with 0.5 mM reduced glutathione in HTF media (Millipore) and spontaneous removal of the cumulus complexes, the number of total and mature oocytes containing single polar bodies at the M2 stage were manually counted in M2 media (Millipore). Oocytes of good morphology lacking polar bodies were defined as M1 stage (immature). Abnormal or fragmented oocytes were defined as oocytes containing multiple cell fragments or multiple polar bodies. Zygotes were rinsed in KSOM media (Millipore) droplets with mineral oil (Sigma) overlay and cultured to blastocyst stage. Ovaries were snap frozen in liquid nitrogen and stored at −80°C for downstream use.

### Histology

Ovaries were preserved in 10% formalin for 20-24 hours and placed in 70% ethanol. Ovaries were then embedded in paraffin and sectioned at 4-5μm per slide. The slides underwent H&E, PAS, and TUNEL assay staining per protocol at Baylor Pathology Core and Lab (H&E, PAS) and Baylor Human Tissue Acquisition & Pathology Core (TUNEL).

### Single nucleus RNA sequencing

Cell nuclei from frozen pooled ovaries (n=4-5) per cohort of young and aged mice from three independent experiments were extracted according to the Chromium Nuclei Isolation Kit (10x Genomics) instructions. Cell nuclei were counted (Logos Biosystems Luna-FL) and visualized under microscopy (Leica DM IL LED) with cell viability <3% with optimal morphology (nuclei with well-resolved edges without debris/clumping). RNA extraction and 10x Single Cell 3’ RNA library preparation v3 were performed according to 10x Genomics manufacturer’s instructions. Our libraries were sequenced using Illumina NovaSeq with paired end 150bp generating average of 376 million reads per sample. About 95% of reads were mapped to the mouse genome. Based on the unique barcodes of each cell, we obtained an average of 46,000 cells per sample totaling 275,781 cells across 6 samples before sample processing. Ambient RNA was removed with SoupX to correct for contamination by cell-free RNA and improve cell-type specific quantification^62^.

### Single cell sequencing analysis

Raw data in FASTQ file format were processed using Cell Ranger 7.0.0 (10x Genomics) with default parameters and cellranger count include-introns for individual samples. Alignment was performed with STAR using reference mouse genome refdata-gex-mm10-5.0.0. Cell calling was performed using the EmptyDrops method in Cell Ranger based on UMI counting. Single cell data processing, clustering, and inferring trajectories was performed according to best practices^63^. Ambient cell-free RNA was removed using SoupX^62^ package with load10x, autoEstCont and adjustCounts functions after basic clustering in Seurat from cellranger output. Corrected counts after SoupX adjustment or raw counts without SoupX adjustment were loaded into scanpy^64^ in a python notebook (v.3.12.4). Cells expressing fewer than 200 genes and genes detected in fewer than 3 cells were filtered out. Cells with mitochondrial counts exceeding 25% were removed, as a relatively permissive threshold previously applied to ovarian single-cell RNA sequencing datasets to retain ovarian cell populations with high mitochondrial content^2^. Doublet detection and removal was performed with scrublet^65^. Counts were normalized by count depth scaling and log1p transformation. The top 2000 genes were annotated as highly variable genes for feature selection. Dimensional reduction was performed by principal component analysis. Samples and percent mitochondrial counts were projected onto UMAP to assess batch effect. Batch correction was applied using harmonypy^66^ for each sample. To generate clusters, we used the Leiden graph-clustering method and re-assessed filtering by visualizing predicted doublets and doublets scores using UMAP to ensure minimal to no batch effect. To optimize cell clustering, we assessed Leiden resolutions of 0.25, 0.50, and 1.00 and compared marker gene expressions of each cluster at 1.00 resolution in UMAP projections. Marker genes were obtained from single cell sequencing data by CellMarker2.0^24^, Mantri^25^, Morris^19^, Russ^26^, Jin^27^, Huang^20^, and Isola^2^ and colleagues. Differentially expressed genes among the clusters were obtained by Wilcoxon and t-tests comparing clusters to one another. The top 100 differentially expressed genes from each cluster were queried in Metascape^67^ for biological functions for annotating clusters. Cell type proportion compositional analysis was conducted using a Bayesian model scCODA^68^ with false discovery rate of 0.05. For dimensionality reduction, we used Potential of Heat-diffusion for Affinity-based Transition Embedding (PHATE)^34^ with following parameters: knn = 30, decay=15.

**Pseudobulk enrichment analysis** was performed by extracting the CL progressing cluster and comparing differential expression between aged vs young (n=3) pooled biological samples using decoupler-py v2.1.6^69^ and pyDESeq2^70^. For the Harmony CL1 and no-SoupX comparisons, differential expression was likewise tested at the biological-sample level. We discarded pseudobulk samples that contained less than 10 cells or fewer than 1000 counts. We used a univariate linear model (ulm) method for enrichment analysis of gene regulatory network CollecTRI, pathway gene sets PROGENy and MSigDB Hallmark datasets.

**Cell to cell communication** was inferred using LIANA+^71^(v1.8.0) with rank_aggregate^72^ method to predict highly ranked ligand-receptor interactions across multiple methods including CellphoneDB, Connectome, log2FC, NATMI, SingleCellSignalR, CellChat, and Geometric Mean. Mouse genes were converted to human gene homologs with pybiomart.

### RNA velocity

Unspliced and spliced read predictions were generated from aligned BAM files using velocyto^73^. The loom files were then combined with annotated anndata files from Cell Ranger. In the python notebook interface, we used scVelo^74^ (v.0.3.4) stochastic, steady-state, and dynamic (EM) models to calculate RNA velocity of cell clusters. Analysis assumed that degradation kinetics represented nuclear export rather than RNA degradation^74^

For sensitivity analysis, using scanpy^64^, we removed filtered outliers using threshold 5 MAD, removing cells with greater than 25% mitochondrial counts^2^. Doublet detection and removal was performed with scrublet. Counts were normalized by count depth scaling and log1p transformation.

### Whole genome bisulfite sequencing (WGBS)

Frozen mouse ovaries were pooled (4-5 mice per experiment and n=3 experiments from young and aged cohorts, respectively). Total DNA was extracted and quantified using Qubit fluorometer by Novogene. DNA integrity for each sample was verified by agarose gel electrophoresis. Library preparation and sequencing were performed at Novogene Corporation Inc. (Sacramento, CA, USA). Briefly, DNA was spiked with lambda DNA, fragmented to 200-400bp and treated with bisulfite to convert unmethylated cytosines to uracil. Illumina adapters were then ligated followed by standard library preparation protocol per Novogene. Libraries were sequenced on Illumina NovaSeq with paired-end 150bp configuration generating 63GB of raw data per sample.

### WGBS data analysis

The quality of raw reads was assessed with FastQC (fastqc_v0.11.8) and trimmed with Trimmomatic v0.36 using the parameters (SLIDINGWINDOW: 4:15); LEADING:3, TRAILING:3; ILLUMINACLIP: adapter.fa:2:30:10; MINLEN:36). The bisulfite conversion rate was noted to be >99.6%. After trimming, we used FastQC to assess the quality of the clean reads. Bismark v0.24.0^75^ was used to align bisulfite-treated reads to mm10 reference genome (-X 700 –dovetail). The reference genome was transformed into the bisulfite-converted version and then indexed using bowtie2^76^. Sequence reads and depth and coverage were summarized using deduplicated reads function. The results of the methylation extractor (Bismark_methylation_extractor, --no_overlap) were transformed into bigwig format for visualization in an IGV browser. The non-conversion rate was defined as the percent cytosine sequenced in the lambda genome. To estimate the methylation site, the sum Mc of methylated counts was modeled as a binomial (Bin) random variable with methylation rate. The bin size was set to 10kb. The sum of methylated and unmethylated read counts in each bin were calculated. The methylation level (ML) for each window or bin or C site is the fraction of methylated Cs and is defined as:

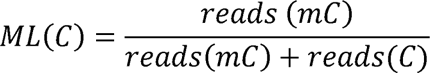

Mean base coverage across samples was 19x. Differentially methylated regions (DMRs) were identified using Dispersion Shrinkage for Sequencing (DSS) method^77–79^. Genes related to DMRs were defined as those whose gene body region (from transcription start site to transcription end site) or promoter region (upstream 2kb from the transcription start site) have an overlap with the DMR.

Enrichment analysis of DMRs was conducted using the Genomic Regions Enrichment of Annotations Tool (GREAT v.4.0.4)^80,81^ using total DMR as background regions, mouse NCBI build 38, basal+extension (constitutive 5.0kb upstream and 1.0kb downstream, up to 1Mb max extension, including curated regulatory domains. Associated annotated genes were analyzed using EnrichR^82–84^.

For promoter regions, gene ontology (GO) enrichment analysis was conducted using GOseq^85^ R package including DMR-related genes with corrected p-value less than 0.05. KEGG^86^ pathways were assessed using KOBAS^87^ software for statistical enrichment of DMR-related genes. The FDR correction method used was Benjamini and Hochberg. Gene bodies were defined as the interval from the transcription start site (TSS) to the transcription end site.

### Reverse phase protein array (RPPA)

Snap frozen ovaries (n=5) in each cohort underwent protein extraction by a core facility at Baylor College of Medicine as previously described^88^. Antibody-based proteomics was performed using 261 previously validated antibodies using an Aushon 2470 robotic protein arrayer (Aushon BioSystems, Billerica, MA) and a high throughput automated immune slide stainer Autolink 48 (Dako, Carpinteria, CA) at the Baylor Proteomics Core Facility^41^. Briefly, samples were affixed onto nitrocellulose-coated slides and incubated with primary antibody for 30min, followed by a secondary antibody with appropriate negative and positive controls. At the time of analysis, we used 261 validated antibodies from the following pathways: DNA damage response (ATM/ATR/CHK/p53), growth factor signaling through PI3K-AKT-mTOR and MAPK, receptor tyrosine kinase, ERK signaling, apoptosis, NF-kB, epithelial-mesenchymal transition, adhesion, invasion, cytoskeleton remodeling, WNT, developmental, stem cell signaling, steroid hormone receptor, epigenetic and chromatin regulation, autophagy and metabolic stress pathways. Signals were amplified with the VECTASTAIN Elite ABC-HRP Kit (Vector Laboratories) and the Tyramine Signal Amplification plus biotin kit (Akoya Biosciences) and detected using fluorescent IRDye 680 streptavidin (Li-Cor Biosciences). Fluorescent signals were recorded at 635nm or 525nm wavelengths with GenePix AL4400 scanner and normalized to total protein detected with Sypro Ruby Blot Stain (Molecular Probes). Signal intensity (SI) was normalized as previously described^41^. Mean normalized SI values were obtained for three technical replicates for each sample and for each protein. Data were analyzed using two-sided t-tests with multiple testing correction using the Storey and Tibshirani^89^ method. Hierarchical clustering was performed using Cluster 3.0^90^. Data was filtered for missing values, standard deviation less than 2.0 with at least one observation with absolute value greater than 2.0, and all genes whose maximum minus minimum values are less than 2.0. Data were log2 transformed, centered on the sample median, and expression was normalized. Normalized expression was visualized with seaborn and matplotlib in python. Differentially expressed signals with greater than an absolute log2-fold change and p<0.05 were analyzed by ingenuity pathway analysis (IPA, Qiagen).

### Immunohistochemistry

Ovarian tissues were submitted to the institutional cores for processing, paraffin embedding, and sectioning. Immunohistochemical staining for CHK2 and phosphorylated CHK2 was performed by the Baylor Pathology Core and Lab. Primary antibodies included CHK2 (Santa Cruz sc-5278, 1:30) and phospho-CHK2 (Thr68) (clone C13C1, Cell Signaling Technology, 2197T). Additional immunohistochemical staining was performed by the Baylor Human Tissue Acquisition & Pathology Core using antibodies γH2AX (phospho-H2AX Ser139, clone JBW301, Millipore 05-636, 1:200 room temperature for 30 minutes), p53bp1 (NB100-304 Novus Biologicals, 1:100, room temperature for 30 minutes), and Rad51 (abcam, ab176458, 1:100, room temperature for 30 minutes). Tissue sections were deparaffinized and re-hydrated, followed by antigen retrieval using antigen retrieval buffer low pH Biocare Diva Decloaker, pressure cooker heated at 110°C for 15min. Then TBS-T wash buffer was used. Images of bright field slides were taken with ImageXpress Pico (Molecular Devices).

### Primary human granulosa cell culture

Discarded follicular fluid was obtained from the Yale Fertility Center after routine oocyte retrieval under approved Yale Institutional Review Board protocol #2000038061. Deidentified information including age and diagnoses were collected to classify samples by age and to exclude patients with polycystic ovarian syndrome. Cells from follicular fluids were separated by centrifugation at 1500rpm for 5 minutes and resuspended in 5ml of ACK lysis buffer (Quality Biological A10492-01) per 50ml volume. Red blood cells were lysed for 5 minutes at 37°C and then washed once with PBS then twice with DMEM/F12 culture media (Invitrogen 11330-032), supplemented with 1% penicillin/streptomycin (Invitrogen 15140-122), 2% glutamax (Invitrogen 35050-061), and 5% fetal bovine serum (Gibco 26140-079). Cells were plated in a T-75 flask overnight and the culture media was changed every two days. On day 4 after isolation, adherent cells were dissociated and replated in 24-well plates. After allowing cells to adhere for approximately 2 hours, the medium was replaced with DMEM/F12 containing 0.1% bovine serum albumin, 1% penicillin/streptomycin, GlutaMAX, and 10^−7 M androstenedione. Cells were maintained under these serum-free conditions for 24 h. Cells were treated with fresh medium containing 0.1% bovine serum albumin and 10^−7 M androstenedione supplemented with follicle-stimulating hormone (FSH 2.5IU) and either vehicle or the CHK2 inhibitor CCT241533 (1μM, MedChemExpress). After 48 h of treatment, cells were harvested for RNA isolation and subsequent gene-expression analysis. Cell culture media was harvested for cytokine analyses.

### RNA analysis via quantitative polymerase chain reaction

Total RNA was isolated from primary human granulosa cells using a column kit (Qiagen RNeasy mini 74104). cDNA was made by reverse transcriptase (Thermo Fisher Scientific 4368814). Equal concentrations of cDNA were used per sample for quantitative PCR together with a SYBR green master mix (MedChemExpress HY-K0501). Fluorescent primer intensities were detected using fluorescent detector (Bio-Rad CFX Maestro Version 4.1.2433.1219). The ΔΔCt normalization method was used with three technical replicates. The primers used for human CD74 were: forward 5’-AAGCCTGTGAGCAAGATGCGCA and reverse 5’- AGCAGGTGCATCACATGGTCCT. The primers used for human IL-1β were: forward 5’- GGATATGGAGCAACAAGTGG and reverse 5’ ATGTACCAGTTGGGGAACTG. The primers used for reference gene GAPDH were: forward 5’-ACA ACT TTG GTA TCG TGG AAG G and reverse 5’- GCC ATC ACG CCA CAG TTT C.

### Bulk RNA sequencing of primary human granulosa cells

Total RNA was isolated from primary human granulosa cells using a column kit (Qiagen RNeasy mini 74104). Total RNA was submitted to Plasmidsaurus for stranded 3’-end counting RNA-seq with unique molecular identifiers (UMI)-based molecule counting using Illumina NovaSeq X Plus sequencing. Reads were quality filtered, aligned to the reference genome, de-duplicated using UMIs. Gene expression was quantified using featureCounts. Differential gene expression analysis was performed using the Plasmidsaurus RNA-seq analysis pipeline. Cells from one donor were allocated untreated, FSH-treated, CCT241533-treated, and FSH plus CCT241533-treated groups with six replicates per group. The total number of reads per sample was 20 million reads except for two samples, which had <19 million reads.

### Immunofluorescent analysis of primary granulosa cells

Cells were cultured on 5mm microscopy glass round sections. On the day of harvest, cells were washed with PBS and were fixed with 4% paraformaldehyde for 30 minutes onto glass cover slips. Then they were resuspended in sterile PBS. Glass cover slips containing cells were transferred to microscope slides and stained with primary and secondary antibodies; and mounted with DAPI by iHisto. Immunofluorescent images of the whole slides were taken with the Pannoramic MIDI II (3DHistech) and viewed with SlideViewer (3DHistech). Images were quantified in a Fiji/ImageJ. Primary antibodies used for immunocytochemistry were γH2AX (phospho-H2AX, Ser139, clone JBW301, MilliporeSigma/EMD Millipore, 05-636), CD3 (nb600-1441ss), CD74 (77274), CD19 (nbp2-25196), CHK2 (Proteintech 13954-1-AP). TUNEL assay staining was performed per protocol by iHisto, Inc. Immunofluorescent images were taken with the Pannoramic MIDI II (3DHistech) and viewed with a SlideViewer (3DHistech).

### Hormone and cytokine assay

For hormone stimulation, serum free DMEM/F12 media was used, supplemented with 0.1% BSA, penicillin/streptomycin, L-glutamine, and 10^−7^ M androstenedione (Steraloids, Inc, A6030-000). The next day, 0.025, 0.25, and 1.25IU/mL of human recombinant follicle stimulating hormone (VWR International 75791-802) were added to the culture media by refreshing the media containing FSH. Following 48 hours of stimulation, culture supernatant was harvested and estradiol concentrations were measured using the estradiol (E2) competitive enzyme immunoassay (Tosoh Bioscience) on an AIA-900 automatic clinical grade automated immunoassay analyzer at Allcare Fertility Center. Samples were analyzed using the manufacturer’s protocol and calibrated according to the manufacturer’s instructions. IL-1β (R&D Systems DY201) was detected in the supernatant by ELISA according to manufacturer’s instructions. Experiments represent data from one donor. Experiments with additional donors were repeated 6 times.

### Human ovarian GTEx data analysis

Expression analysis of CHEK2 was obtained from Adult Genotype-Tissue Expression (GTEx) Portal^91–93^. Bulk tissue gene expression for *CHEK2* (ENSG00000183765.23) was obtained via GTEx Analysis Release V10 (dbGaP Accession phs000424.v10.p2) on 01/09/2026. Data were filtered to include a maximum of two tissue types per organ. Cell lines were removed and transcripts per million (TPM+1) values were log-transformed. Normalized TPM for RNA-seq data were obtained from the Human Protein Atlas Tissue Ovary-GTEx RNA-seq data set. Mean nTPM values were plotted and statistical testing was performed using Graphpad Prism 10.

### Gene set enrichment and rank-rank hypergeometric overlap analysis

Preranked gene set enrichment analysis (GSEA)^94^ was performed independently for human granulosa cells treated with CHK2 inhibitor plus FSH versus FSH alone and for aged versus young mouse antral granulosa cell clusters using gseapy^95^. Enrichment was tested against the MSigDB Hallmark 2020^96^ gene sets, and pathways with FDR *q* < 0.05 were considered significant. To compare the human granulosa cell CHK2-inhibition and mouse ovarian-aging transcriptional responses, we performed rank-rank hypergeometric overlap (RRHO) analysis^49^. Human genes were mapped to mouse orthologs using gProfiler^97^, and analysis was filtered for unique genes shared between the two datasets. The complete ortholog-matched gene lists were ranked independently according to the direction and strength of differential expression. Overlap across pairs of rank thresholds was evaluated using a hypergeometric distribution. For directional analysis, each ranked profile was evaluated from both ends of the distribution, generating four combinations: CHK2i-up × Age-down and CHK2i-down × Age-up (transcriptional reversal), and CHK2i-up × Age-up and CHK2i-down × Age-down (transcriptional concordance). Seventy-five rank thresholds spanning each directional gene list were evaluated. At each pair of thresholds, enrichment of overlapping genes relative to the complete shared ortholog universe was quantified using a one-sided hypergeometric test. Thresholds were expressed as the percentage of the ranked gene universe included at each position. For visualization, the four directional RRHO surfaces were combined into a single four-quadrant map. Hypergeometric significance was represented as signed −log10(*P*), with positive values assigned to the two reversal quadrants and negative values assigned to the two concordant quadrants. Thus, the magnitude indicates the strength of overlap enrichment, whereas the sign denotes reversal or concordance. The individual hypergeometric *P* values in the RRHO surface were used to visualize local enrichment.

## Manuscript editing

We thank Angelika H. Hofmann at SciWri Services for manuscript editing. ChatGPT was used to improve the readability of the manuscript.

## Acknowledgements

We thank Xiaojun Xing and Tim Nottoli at the Yale Genome Editing Center (YGEC), Cihan Halicigil, Stephanie Nichols-Burns, Olga Chaplia, and Murat Basar at Yale Fertility Center, Mingxue Yang and Yula Ma at Allcare Fertility, and members of the Heaney, Lee, and Jiang laboratories for technical assistance. LY is supported by the American Society for Cell and Gene Therapy Career Development Award 2022, the Yale Women’s Reproductive Health Research Career Development Center (Eunice Kennedy Shriver National Institute of Child Health & Human Development of the National Institutes of Health under Award Number K12HD047018), Reproductive Scientist Development Program (K12HD000849), Albert McKern Award, Bennack-Polan Foundation Award. LY is a Pepper Scholar with support from the Claude D. Pepper Older Americans Independence Center at Yale School of Medicine (P30AG021342). JDH is supported by the BCM Knockout Mouse Production and Phenotyping Project UM1 HG006348 and BCM Center for Precision Medicine Models U54 OD030165. Baylor Tissue Acquisition & Pathology (HTAP) and the Baylor Genetically Engineered Rodent Models Core are supported by the Dan L. Duncan Comprehensive Cancer Center grant P30 CA125123. Baylor RPPA Core was supported by Cancer Prevention & Research Institute of Texas Core Facility Support Awards (PR210227, PI: Dean Edwards). The content is solely the responsibility of the authors and does not necessarily represent the official views of the National Institutes of Health. Graphical images were made using BioRender.com.

## Author Contributions

LY, MT, HST, YJ, BL, TPW, and JDH designed the study.

LY, ZS, IL, AP, and MD conducted the animal experiments.

LY conducted the ovarian cell experiments and analyzed bulk RNAseq data.

LY and XL interpreted the WGBS data and analyzed snRNAseq data.

ZS conducted cryosectioning and immunofluorescence experiments and analyses.

CL processed the immunohistochemistry imaging and conducted analysis.

HST, YJ, BL, TPW, JDH supervised the overall project.

LY, MT, HST, YJ, BL, TPW, and JDH interpreted the results, and LY wrote the paper with input from all authors.

## Competing interests

The authors have no conflicts to declare.

## Data availability

The sequencing data and expression count matrices have been deposited in the Gene Expression Omnibus (GEO) and will be made publicly available upon publication.

## Code availability

Code used by the authors is available at https://github.com/LiubinYangResearch/MultiOmic_ovarian_aging

## Extended Data

**Table 1. Annotation of ovarian cell clusters identified by single-nucleus RNA sequencing.** Cell-type annotations for 14 clusters assigned using differentially expressed genes, biological functions and pathways, and comparison with published ovarian cell populations and marker genes. The table includes proposed cell identities, annotations from reference studies, predicted regulators, and tissue annnotations.

**Table 2. Marker genes for each ovarian cell cluster** The table contains 100 ranked genes for each of 14 clusters (groups 0-13) with gene names, differential expression scores, log fold changes, nominal P-values, and multiple-testing adjusted P values.

**Table 3. Pseudobulk differential expression results for aged versus young CL progressing cluster.** Full pyDESeq2 results for 26,673 genes in the CL progressing cluster, using biological samples as independent observations (three per age group). Columns report gene symbol, baseMean (mean normalized count across samples), log2FoldChange (aged relative to young), lfcSE (standard error of the log2 fold change), stat (Wald statistic), pvalue (nominal P value), and padj (multiple-testing-adjusted P value). The table includes all tested genes; blank entries denote unavailable estimates or test results.

**Table 4. Whole-genome bisulfite sequencing quality control results.**

**Sheet 1**. Table of overview of mapping results per sample including total reads denoting filtered clean reads used for mapping, mapped reads is uniquely mapped reads, mapping rate is percentage of uniquely mapped reads in the filtered reads, and duplication rate is the percentage of duplicated reads in the filtered reads. **Sheet 2.** Table of genome coverage for each sample. Coverage is calculated the total reads covering each base

**Table 5. Differentially methylated regions between young and aged ovaries.** Differentially methylated regions (DMRs) identified by whole-genome bisulfite sequencing of aged and young ovaries are listed with their genomic coordinates, region length, number of cytosines analyzed, mean methylation levels in aged and young samples, methylation difference, DMR test statistic, and cytosine context. Methylation differences are aged minus young; positive values denote hypermethylation and negative values denote hypomethylation in aged ovaries.

**Table 6. Normalized RPPA dataset from young and aged ovaries.** LY_1 to LY_5 correspond to young samples, LY_6 to LY_10 correspond to aged samples, LY_11 to LY_15 correspond to aged samples treated with CCT241533.

**Table 7. Differential expression results for primary human granulosa cells treated with FSH and CHK2 inhibitor versus FSH alone.** Results include gene identifiers, log fold change, log counts per million, nominal P value, false discovery rate, test statistic, group-normalized counts and standard deviations, and gene annotations.

**Extended Data Figure 1: Single nuclei RNA sequencing**

**a**, Workflow for annotating ovarian cell clusters. **b**, List of number of cells per annotated cluster grouped by age. **c**, Dotplot of cell-type marker expression by cluster. Color represents mean expression and dot size represents the percentage of cells expressing each marker. **d**, PROGENy pathway activity inferred using a univariate linear model. Positive and negative scores indicate higher and lower inferred activity, respectively, in aged versus young CL progressing cluster. **e**, JAK-STAT geneset enrichment plot. Green line shows the running enrichment score, vertical bars show pathway gene positions. Lower panel shows the ranked-list metric. **f**, UMAP of Harmony-integrated samples colored by sample identity. **g**, UMAP of the same analysis colored by annotated cell cluster. **h**, Boxplots of the proportion of each cell type in a Bayesian model of compositional analysis (scCODA) in the Harmony-integrated dataset. *credible compositional change at a false discovery rate <0.05. **i,** Volcano plot of aged-versus-young differential expression in the Harmony-defined CL1 cluster. Red and blue points indicate increased and decreased expression, respectively; gray points denote genes outside the plotted thresholds. The horizontal axis shows log fold change and the vertical axis shows −log10(P).

**Extended Data Figure 2: Luteal immune signatures and inferred cellular trajectories**

**a**, IPA network depicting predicted activation of HFE in aged luteal cells. Red and blue nodes indicate increased and decreased expression, respectively; edge colors and symbols indicate the IPA relationship predictions. **b**, Predicted upstream regulators from IPA analysis. **c,** CD74 immunohistochemistry in a human ovary from a 33-year-old donor, obtained from the Human Protein Atlas. **d,** CD3 and FOXL2 immunofluorescence in young and aged mouse ovaries, with DAPI nuclear staining and quantification of T cells per follicle from two independent experiments. Mann–Whitney test, ****P<0.0001. **e,** scCODA analysis of cell-type proportions without ambient RNA removal; asterisks indicate credible compositional changes at a target false discovery rate of 0.05. **f,** Volcano plot of aged-versus-young differential expression in the CL cluster without ambient RNA removal. Red denotes increased expression and blue denotes decreased expression. **g,** CollecTRI transcription-factor activity inferred using a univariate linear model. Positive scores indicate greater inferred activity in aged cells and negative scores indicate lower inferred activity. **h,** Metascape enrichment of genes upregulated in aged cells, colored by P value. **i,** IPA graphical summary of predicted biological relationships. **j,** Scatter plot of velocity pseudotime based on the directed velocity graph projected onto the UMAP. **k,l**, Predicted transitions from selected cell populations, colored by the inferred number of steps from the root cell. **m,** PHATE embedding colored by annotated cell type; proximity reflects similarity in gene-expression profiles.

**Extended Data Figure 3: DNA methylation across genomic features**

**a**, Heatmap of CG methylation across genomic annotations in individual young and aged samples. Promoters defined as 2kb region upstream of the transcription start site. Untranslated regions, exons, and introns follow Ensembl annotations. **b,** Methylation profiles across functional genomic regions in young and aged ovaries. Color denotes age group and line style denotes CG, CHG, or CHH context. The left axis shows CG methylation and the right axis shows non-CG methylation (CHG and CHH). **c,** Methylation profiles from 2 kb upstream of the transcription start site through the gene body to 2 kb downstream of the transcription end site. **d,** Circos plot of chromosomal methylation patterns, showing young methylation levels, aged-minus-young methylation differences, and aged methylation levels from outside to inside. **e,** Overlap of DMR-associated gene sets among CG, CHG, and CHH contexts. **f,** Distribution of CG DMR methylation levels in aged and young ovaries. **g,** Distribution of mean methylation levels among DMRs, grouped by age. **h–j**, Numbers of hypermethylated and hypomethylated DMRs across genomic annotations for CG (**h**), CHG (**i**), and CHH (**j**) contexts.

**Extended Data Figure 4: Functional enrichment of differentially methylated regions**

**a,** Gene Ontology (GO) enrichment for genes associated with hypermethylated CG regions. Biological process (green), cellular component (orange), and molecular function (blue); asterisks mark terms with corrected P<0.05. **b,** KEGG enrichment for genes associated with hypermethylated CG regions. **c,** KEGG enrichment for genes associated with hypomethylated CG regions. In b and c, the horizontal axis shows the rich factor, color denotes the q value, and dot size denotes the number of genes. **d,** GO enrichment for genes associated with hypermethylated CG promoter regions. CC, cellular component; MF, molecular function. **e,** GO biological-process enrichment for age-acquired methylated regions generated from GREAT.

**Extended Data Figure 5: Proteomics pathways and DNA damage markers**

**a**, Bar plot of enriched pathways generated in EnrichR in ARCHS4 Tissues gene set of differentially expressed proteins between ovaries of aged and young mice. **b**, Bubble plot of disease and biological functions associated with differential protein expression of aged vs young ovaries using Ingenuity Pathway Analysis Disease and Functions module. Bubbles represent enriched biological function or disease annotation. Positive activation z scores (orange) indicate predicted activation in aged cells and negative (blue) indicate predicted inhibition. Selected highly significant or strongly predicted annotations are labeled. **c**, Immunohistochemistry of representative whole ovary section of young and aged samples. i) regions with corpus luteum stained with TUNEL. ii) regions with antral follicles stained with TUNEL. iii) ovaries stained with anti-RAD51 antibody iv) ovary stained with anti-p53BP1. **d**, Violin plot of log transcripts per million (TPM) of *CHEK2* gene expression from bulk RNA sequencing data from GTEx database, ranked by expression level. **e**, Immunofluorescent images of mouse embryos generated from oocytes of young and aged female mice (1 cohort) stained with anti-p53 antibody with quantification per embryo (right). n=15 embryos per group. HEK293T annotated with arrow. Scale bars indicate standard deviation. **f**, Canonical pathways related to DNA damage and immune activation in differentially expressed genes of CL progressing cluster in pseudo bulk RNA sequencing analysis.

**Extended Data Figure 6: Responses to transient CHK2 inhibition**

**a**, Bar graph of quantification of percentage of mature metaphase II (M2) oocytes in the NIA C57/BL/6 cohort after superovulation. Bars show means with standard deviations. ns is non-significant, *p<0.05. Dots represent each experimental cohort. n=5 animals per cohort over two independent experiments. **b**, IPA canonical pathways associated with differential protein and phosphoprotein abundance in CHK2i-treated versus untreated aged mouse ovaries, measured by RPPA. Bar length indicates −log10(P), and color indicates predicted activation, inhibition, or no directional prediction. **c**, Bar graph of estradiol concentration in supernatant of primary granulosa cells after stimulation with FSH. Mann-Whitney test. Error bar indicates standard deviation. *p<0.05. **d**, MSigDB Hallmark gene-set enrichment comparing CHK2i alone versus untreated (negative-control) primary human granulosa cells. Pathways with FDR <0.05 are shown. Position and color indicate NES. **e**, IPA canonical pathways associated with differential gene expression in CHK2i-treated versus untreated primary human granulosa cells. **f**, Predicted upstream regulators for the same comparison. In e and f, bar length indicates −log10(P), and color indicates predicted activation, inhibition, or no directional prediction. **g,** GO biological-process enrichment for age-upregulated genes suppressed by CHK2i (left) and age-downregulated genes restored by CHK2i (right), using ortholog-matched human treatment and mouse aging responses. The horizontal axis shows signed −log10(FDR); dot size indicates odds ratio. Filled circles denote FDR<0.05 and open circles denote FDR≥0.05.

## Notes

### Competing Interest Statement

The authors have declared no competing interest.

