## Supplemental Figures for "Multi-omic profiling reveals impaired luteal resolution and CHK2-regulated inflammation in ovarian aging"

### Extended Data Fig 1

a

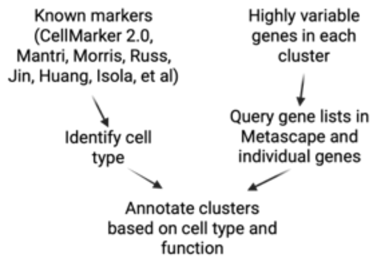


b

| age_group | Aged | Young |
|---|---|---|
| CL regressing | 42948 | 34564 |
| CL mural | 1218 | 2489 |
| CL progressing | 59904 | 298 |
| Stroma | 5084 | 4017 |
| Epithelial | 4779 | 1976 |
| CL ovulatory | 3115 | 11340 |
| Granulosa antral | 17968 | 12989 |
| Granulosa MAPK | 149 | 12333 |
| Endothelial | 4307 | 2859 |
| Theca | 9718 | 3944 |
| Granulosa signaling | 2560 | 7128 |
| Granulosa ECM | 4064 | 5478 |
| Erythrocyte | 1569 | 140 |
| CL regressing steroidogenesis | 13932 | 4482 |
| Total | 171315 | 104037 |

c

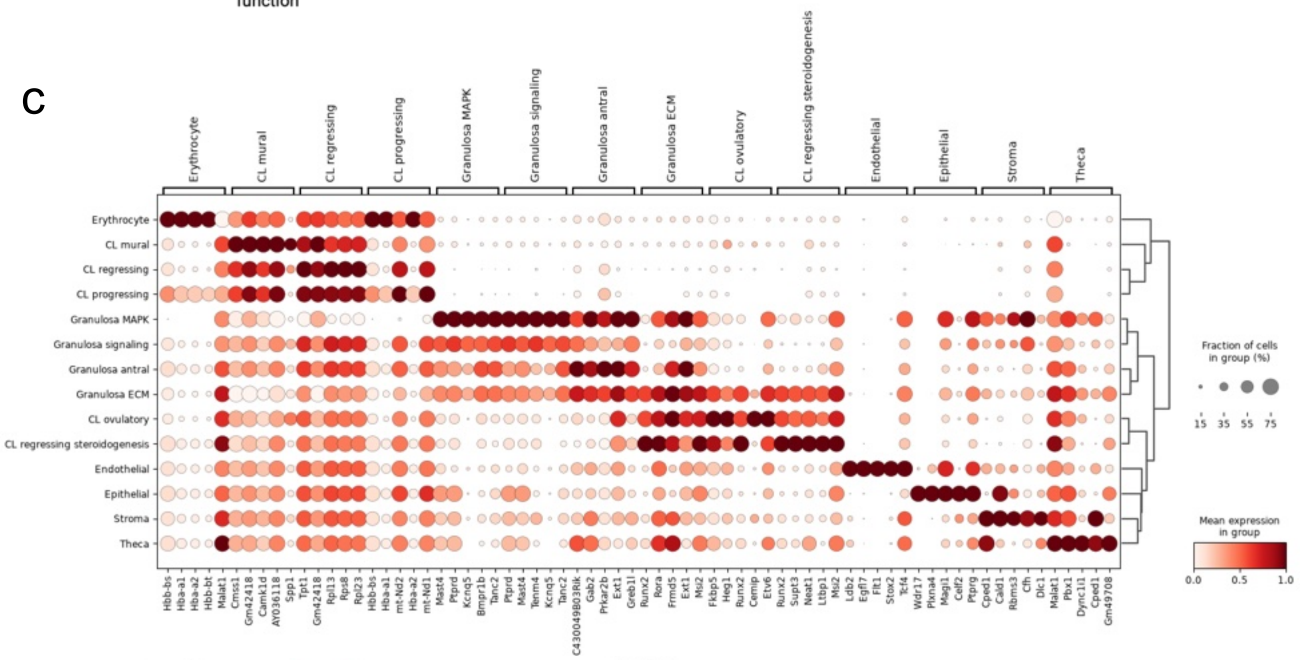


d

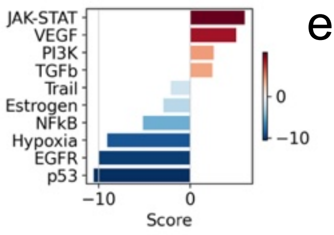


e

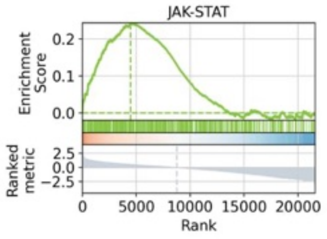


f

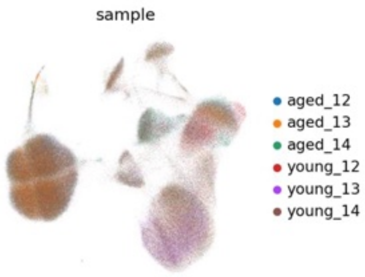


g

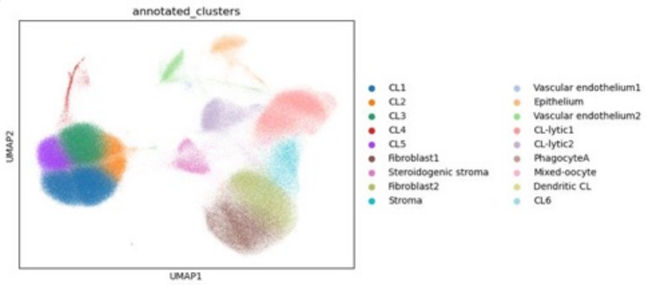


h

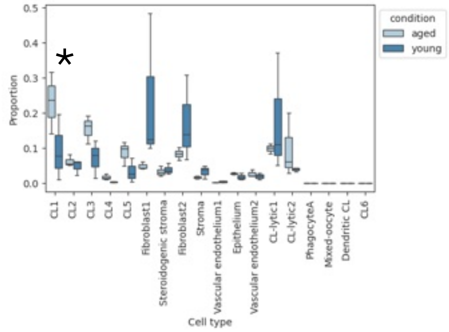


i

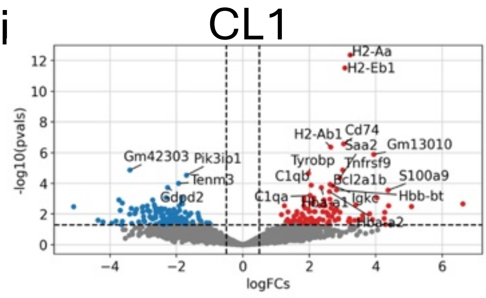

### Extended Data Fig 2

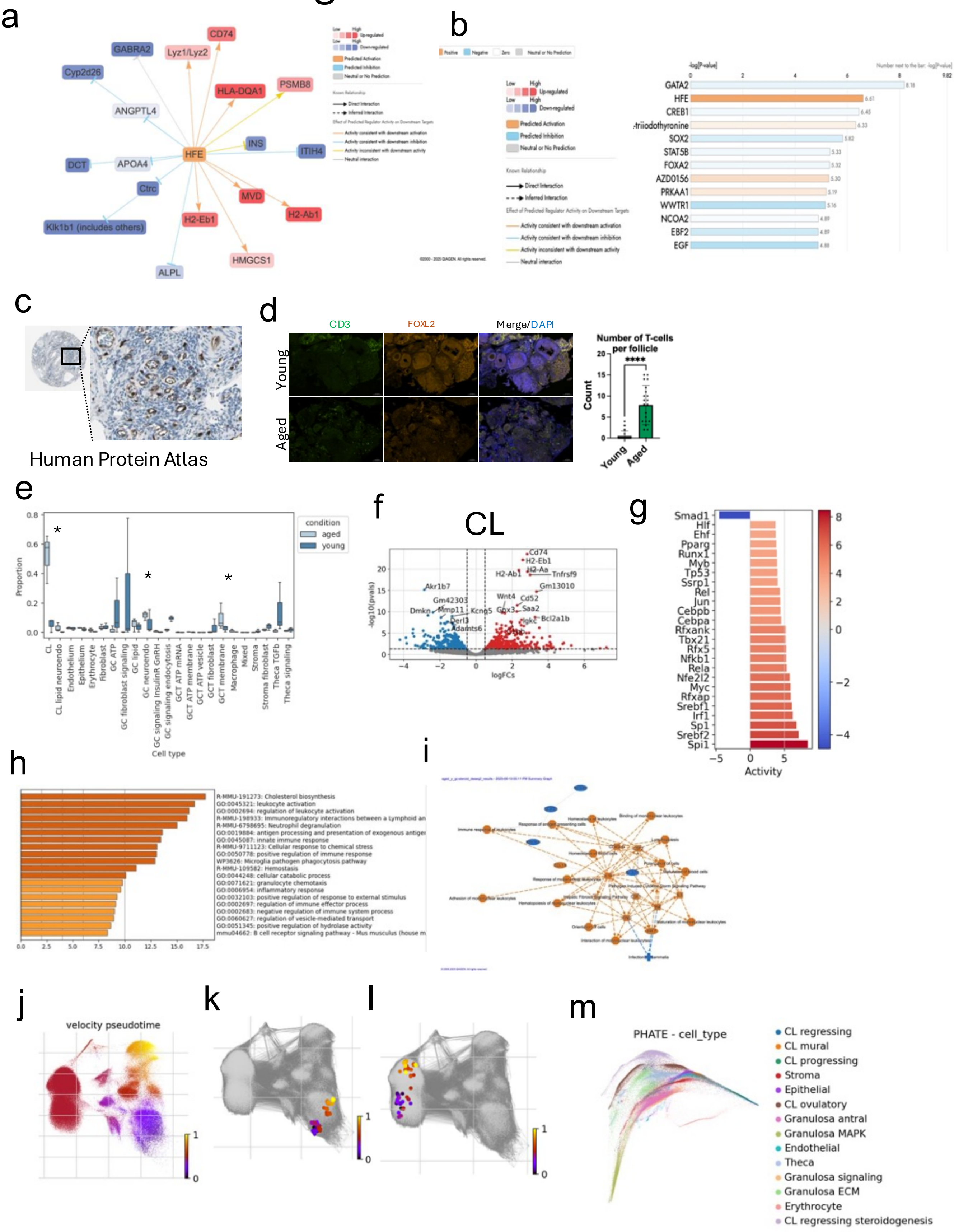

### Extended Data Fig 3

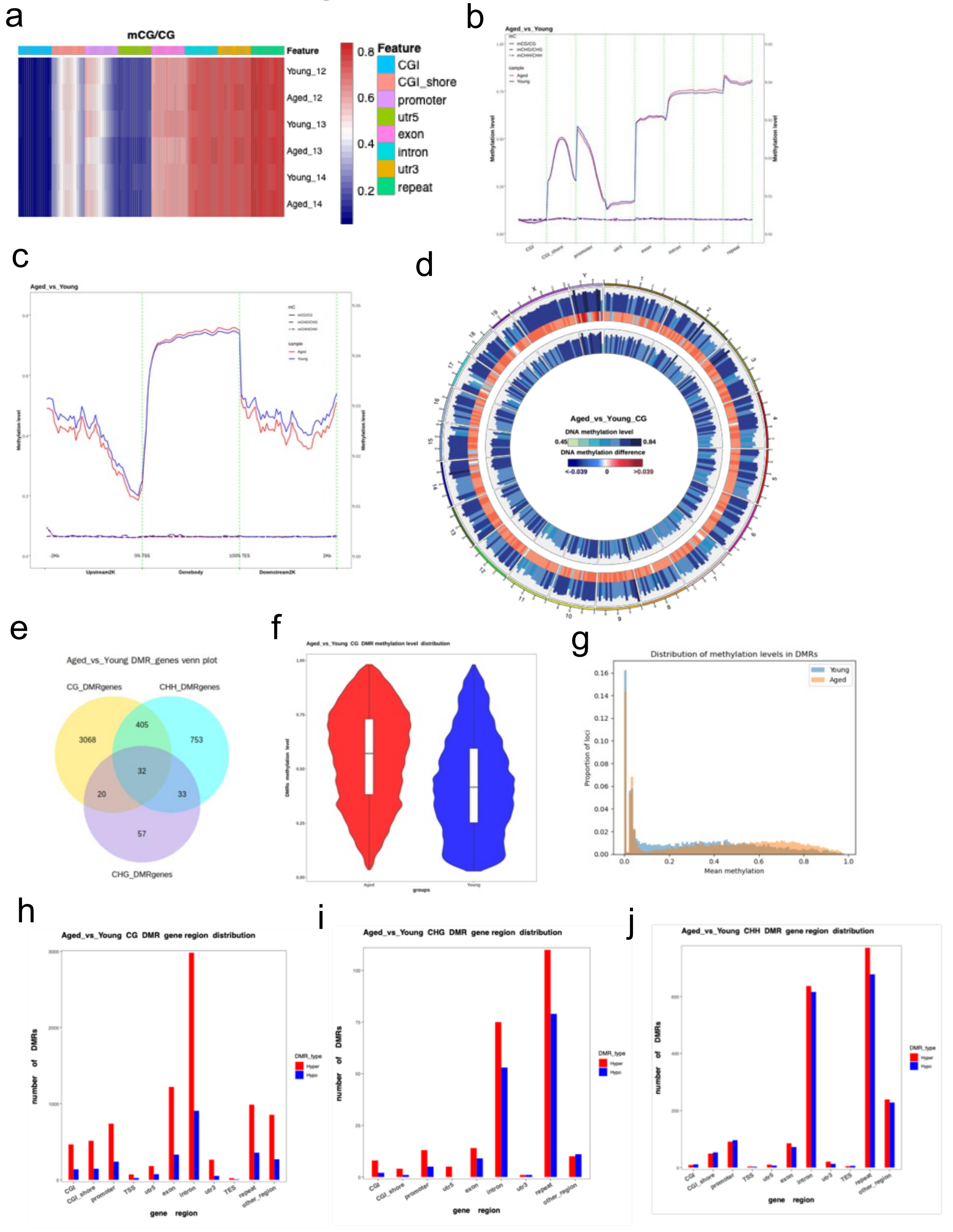

### Extended Data Fig 4

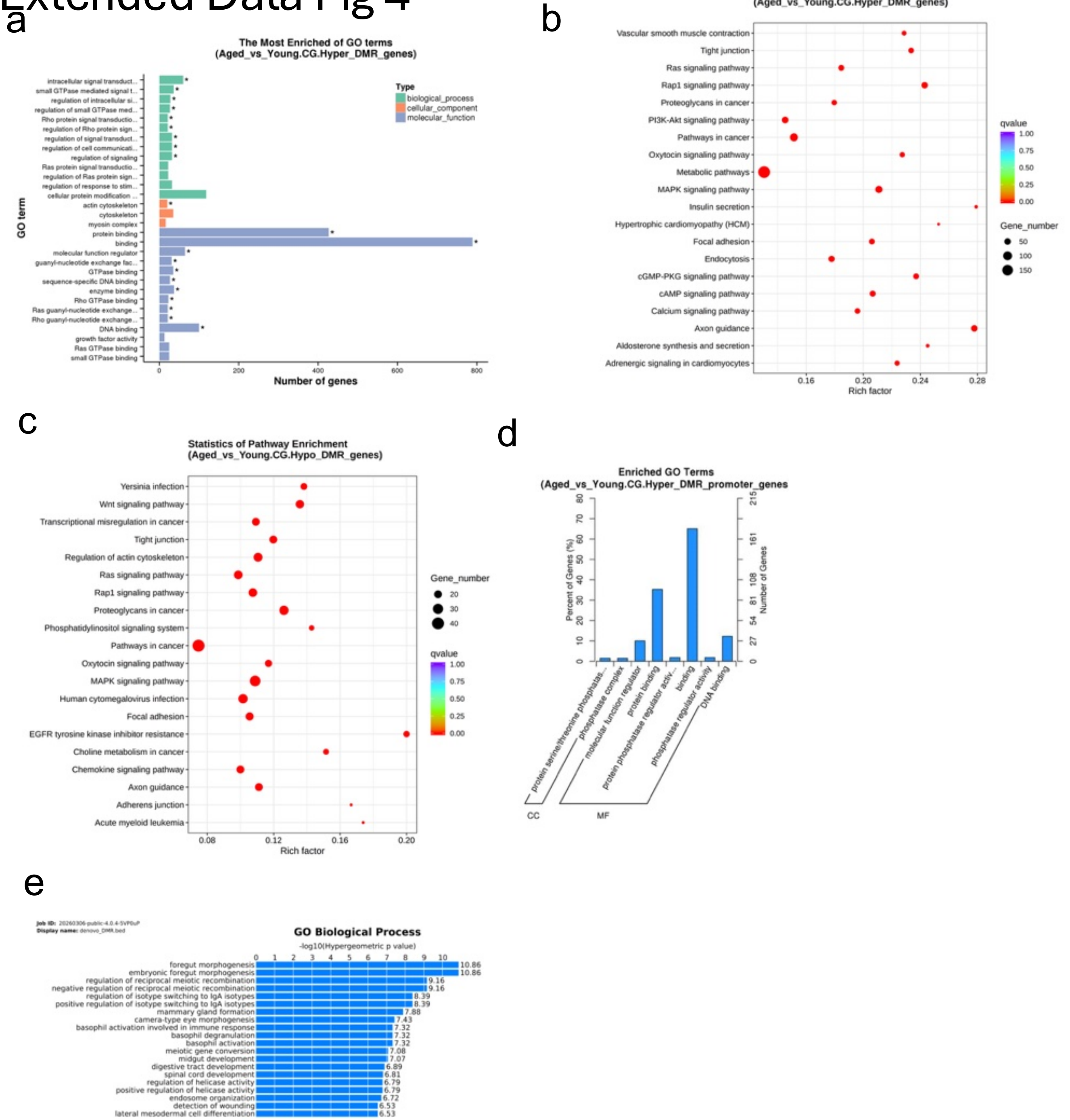

### Extended data figure 5

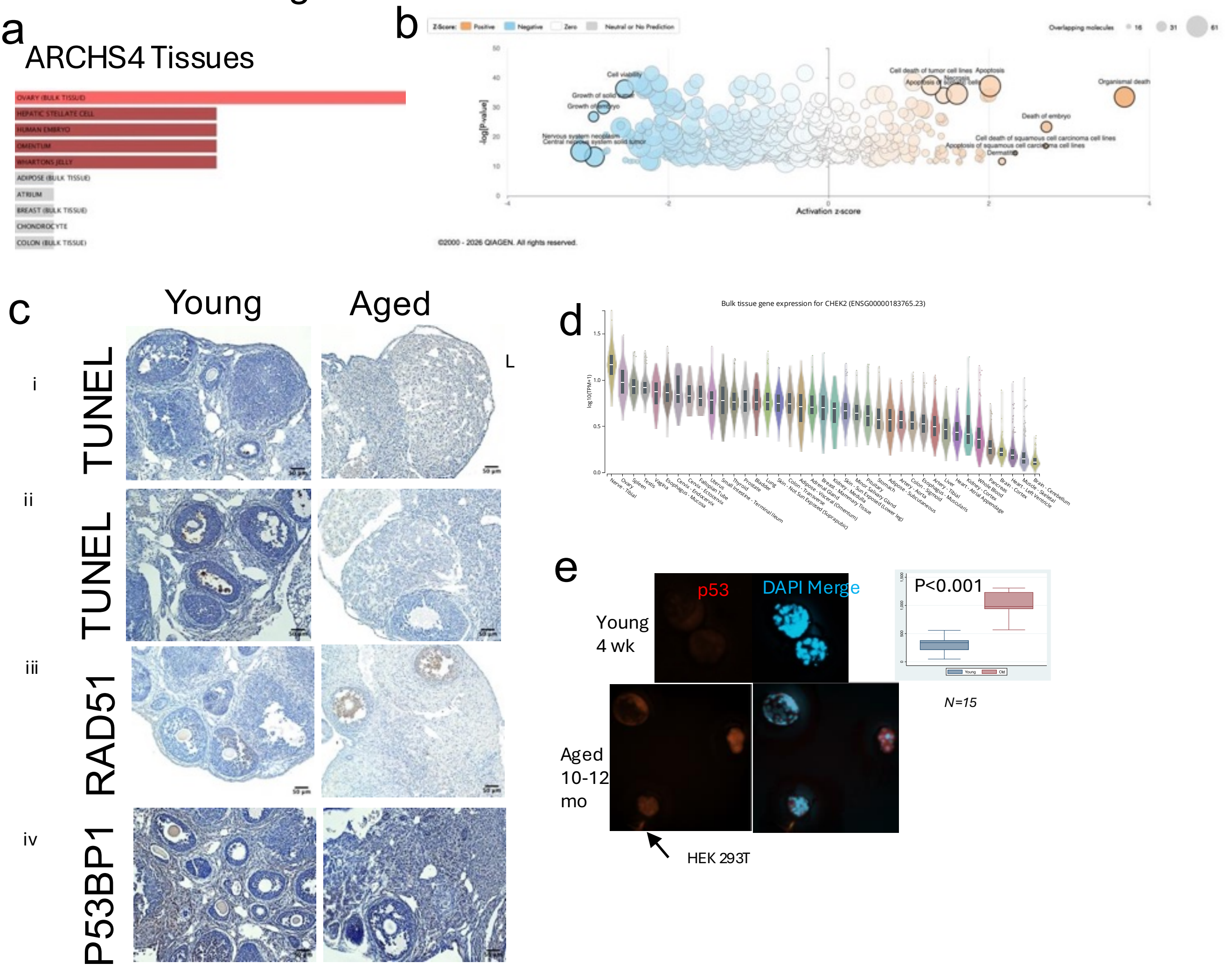


f

| Ingenuity Canonical Pathways | -log(p-value) | Ratio | z-score | Molecules |
|---|---|---|---|---|
| **CGAS-STING Signaling Pathway** | 3.91E-01 | 4.76E-02 | -1.633 | IL12B,IL17B,TNFSF14,TNFSF15,TNFSF4,TREX1 |
| **ATM Signaling** | 0E+00 | 1.02E-02 | NaN | CCNB3 |
| **NFKBIE Signaling Pathway** | 0E+00 | 1.54E-02 | -1.000 | FCER1G,TNFSF14,TNFSF15,TNFSF4 |

### Extended data figure 6

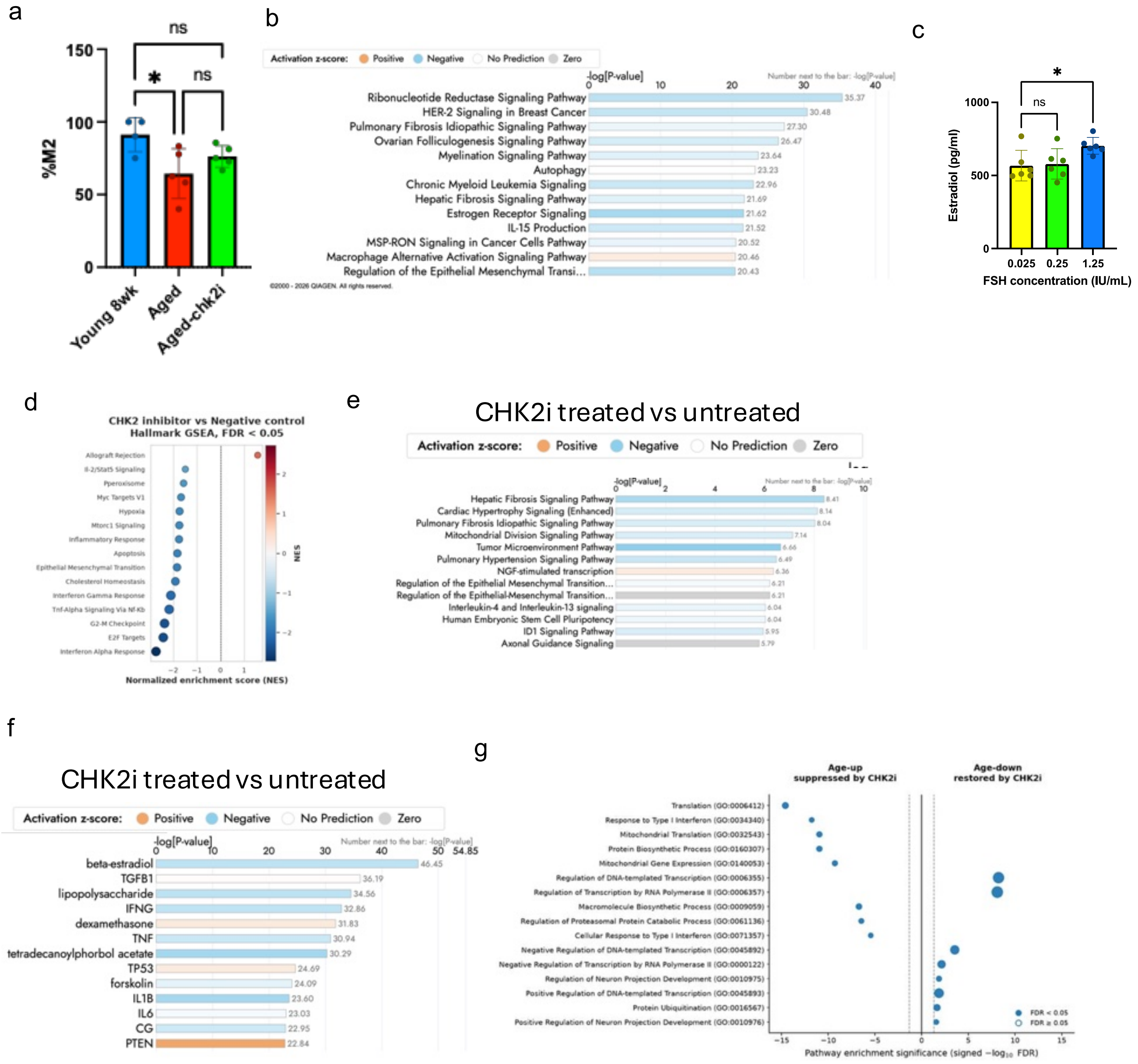
